# Neuronal SEL1L-HRD1 ERAD regulates one-carbon metabolism and is essential for motor function and survival

**DOI:** 10.1101/2025.06.16.659938

**Authors:** Mauricio Torres, You Lu, Brent Pederson, Hui Wang, Anna Gretzinger, Liangguang Leo Lin, Jiwon Hwang, Alan Rupp, Abigail Tomlinson, Andrew J. Scott, Zhen Zhao, Daniel R. Wahl, Martin Myers, Costas A. Lyssiotis, Ling Qi

## Abstract

Hypomorphic variants in the SEL1L-HRD1 ER-associated degradation (ERAD) complex have been linked to severe neurological syndromes in children, including neurodevelopmental delay, intellectual disability, motor dysfunction, and early death. Despite this association, its physiological importance and underlying mechanisms in neurons remain poorly understood. Here, we show that neuronal SEL1L-HRD1 ERAD is essential for maintaining one-carbon metabolism, motor function, and overall viability. Neuron-specific deletion of *Sel1L* in mice (*Sel1L^SynCre^*) resulted in growth retardation, severe motor impairments, and early mortality by 9 weeks of age—mirroring core clinical features observed in affected patients—despite preserved neuronal numbers and only modest ER stress. Multi-omics analyses, including single-nucleus RNA sequencing and metabolomics, revealed significant dysregulation of one-carbon metabolism in ERAD-deficient brains. This included activation of the serine, folate, and methionine pathways, accompanied by elevated levels of S-adenosylmethionine and related metabolites, likely resulted from induction of the integrated stress response (ISR). Together, these findings uncover a previously unappreciated role for neuronal SEL1L-HRD1 ERAD in coordinating ER protein quality control with metabolic adaptation, providing new insight into the molecular basis of ERAD-related neurodevelopmental disease.

**Summary:** Using a neuron-specific *Sel1L* knockout mouse model, we demonstrate that *Sel1L* deficiency activates integrated stress responses, rewires one-carbon metabolism, and impairs motor function and survival.

## INTRODUCTION

Protein synthesis and folding within the endoplasmic reticulum (ER) are tightly coordinated processes essential for cellular homeostasis (1–3). To maintain proteostasis and prevent the accumulation of aberrant proteins, the ER relies on quality control mechanisms, most notably the unfolded protein response (UPR) and ER-associated degradation (ERAD) pathways (4). ERAD targets misfolded or unassembled proteins for retrotranslocation to the cytosol and subsequent proteasomal degradation (5–8). Among ERAD pathways, the SEL1L-HRD1 complex is the most evolutionarily conserved, from yeast to mammals (9–11). HRD1, a multi-pass transmembrane E3 ubiquitin ligase, forms the retrotranslocation channel (8, 9, 12), while its adaptor, SEL1L, ensures HRD1 stability and substrate recognition (13–18). Deletion of Sel1L or Hrd1—either globally or in an inducible fashion—results in embryonic or early postnatal lethality, underscoring their essential roles in development and tissue homeostasis (16, 19–21). Cell type–specific knockout studies have further revealed the importance of SEL1L-HRD1 ERAD in diverse physiological contexts, including lipid and glucose metabolism, immune regulation, thermogenesis, and organ integrity (6, 7, 22–25). Recent work from our group has identified pathogenic variants in SEL1L and HRD1 in pediatric patients presenting with neurodevelopmental delay, intellectual disability, motor dysfunction, and early mortality—clinical features that define a novel syndrome we refer to as ERAD-associated neurodevelopmental disorder with onset in infancy (ENDI) (26, 27).

Neurons, as highly specialized and long-lived postmitotic cells, are particularly dependent on efficient protein folding and ER quality control for lifelong function. The ER is central to the folding and maturation of numerous membrane and secretory proteins vital for neurotransmission, synaptic plasticity, and cellular viability (28, 29). Despite the importance of proteostasis in neurons, the functional role of ERAD in the nervous system remains largely unclear. Elucidating ERAD’s role in neurons is critical not only for understanding the pathogenesis of ENDI but also for exploring broader implications in neurodegenerative diseases and age-associated cognitive decline. Prior studies have shown that deletion of Sel1L in hypothalamic neurons disrupts the maturation of prohormones and receptors—including pro– arginine vasopressin (proAVP), pro-opiomelanocortin (POMC), and the leptin receptor—leading to altered fluid balance and energy homeostasis (30–33). Similarly, selective loss of Sel1L in cerebellar Purkinje cells causes early-onset ataxia and neuron degeneration, supporting a cell-autonomous role for ERAD in neuronal integrity (34). However, the global consequences of neuronal ERAD deficiency in vivo remain poorly defined.

In this study, we generated neuron-specific conditional knockout models targeting SEL1L or the UPR sensor IRE1α to dissect the importance of SEL1L-HRD1 ERAD to brain function. We found that deletion of Sel1L, but not Ire1α, in a subset of neurons led to growth retardation, motor deficits, and early mortality. Remarkably, these phenotypes occurred in the absence of overt neuronal loss or robust UPR activation. Multi-omics analyses, including single-nucleus RNA sequencing and metabolomics, revealed broad upregulation of one-carbon metabolism, a pathway central to nucleotide biosynthesis and methylation reactions essential for brain development (35). Disruption of one-carbon metabolism has been implicated in multiple neurodevelopmental and neuropsychiatric disorders, including neural tube defects, autism spectrum disorder, and schizophrenia, often through impaired methylation, altered neurotransmitter synthesis, and elevated homocysteine levels (35–37). Furthermore, the observed metabolic shifts were accompanied by activation of the integrated stress response (ISR) (38), suggesting that ERAD deficiency leads to metabolic reprogramming rather than classical neurodegeneration.

## RESULTS

### SEL1L and HRD1 are highly enriched in neurons

To assess the cell type-specific distribution of ERAD components in the brain—including neurons and glial cells such as astrocytes, oligodendrocytes, and microglia (39)—we examined SEL1L and HRD1 protein expression in coronal sections from 5-week-old wild-type (WT) mice by immunofluorescence (Figure 1 and Supplemental Figure 1). Neurons were identified by NeuN, a nuclear and perinuclear marker of mature neurons, while astrocytes, the most abundant glia cell in brain, were labeled with glial fibrillary acidic protein (GFAP). In the cerebral cortex, SEL1L was predominantly localized to the perinuclear region of NeuN-positive neurons, consistent with its localization to the ER (Figure 1A-B, D). In contrast, astrocytes – distinguished by their smaller, stellate morphology – exhibited relatively weak SEL1L expression (Figure 1A-B, D, J). In the hippocampus, SEL1L was strongly expressed in neuronal layers of Cornu Ammonis (CA) 1 and CA3, as well as the dentate gyrus (DG), whereas astrocytes exhibited markedly weaker signals (Figure 1C, E-F, H, K-L). Similarly, in the cerebellum, SEL1L expression was notably higher in Purkinje neurons compared to surrounding glial cells (Figure 1G, I, M). HRD1 exhibited a comparable distribution pattern and subcellular localization across these brain regions (Supplemental Figure 1). Collectively, these data demonstrate that the SEL1L-HRD1 ERAD complex is highly enriched in neurons across multiple brain regions, supporting a potential role in neuronal function and maintenance.

**Figure 1.**
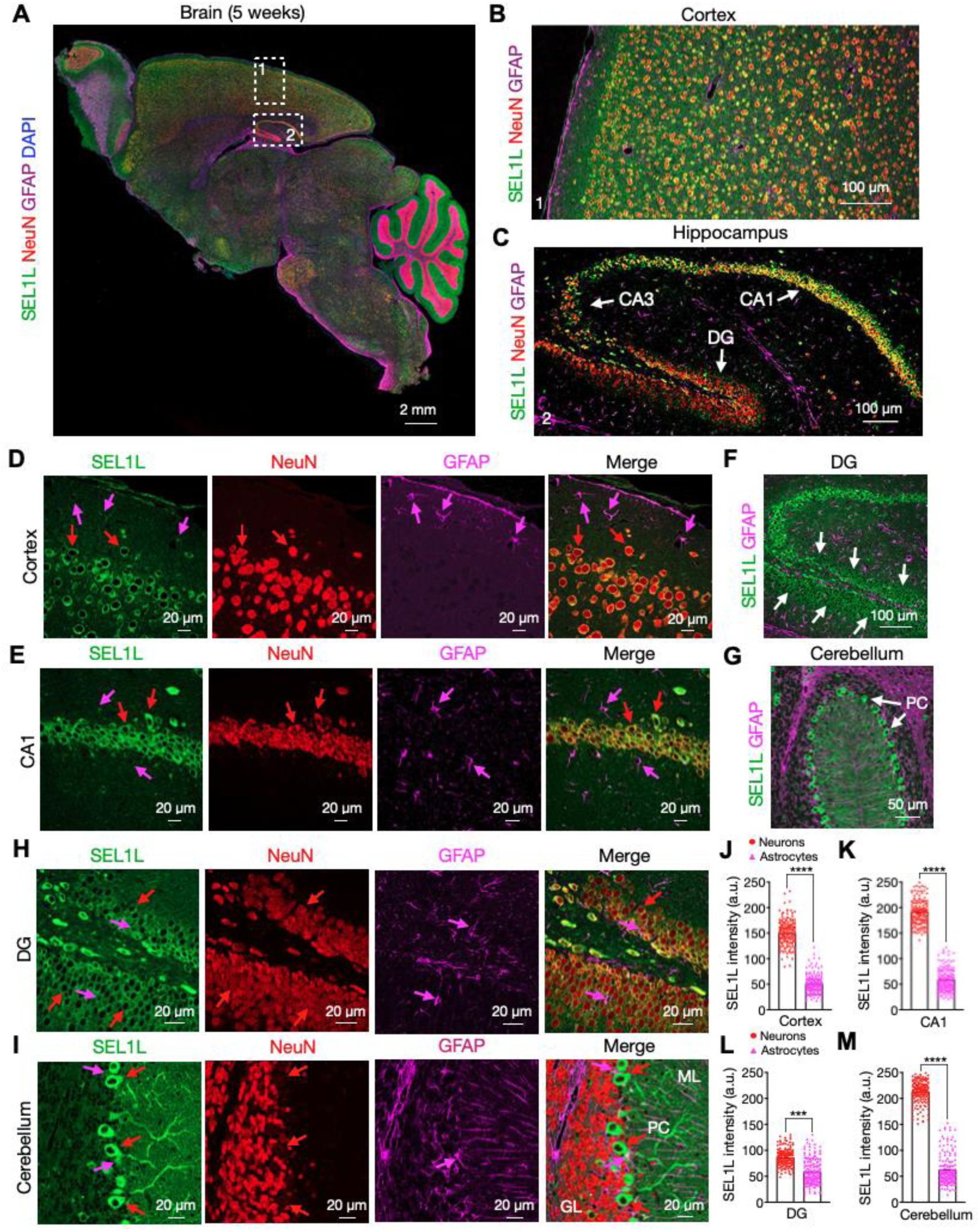
SEL1L protein is highly expressed in neurons. (**A**) Representative confocal image of SEL1L (green), NeuN (red), GFAP (magenta), and DAPI (blue) staining in a sagittal brain section of a 5-week-old mouse. The magnification of the selected regions for cortex and hippocampus, are showed in panels (**B)** and (**C)**, respectively. White arrows in panel (**C**) indicate the CA1, CA3, and DG regions of the hippocampus. (**D** and **E**) Representative zoomed-in images for cortex (**D**) and CA1 (**E**) regions showing separated channels for SEL1L, NeuN, and GFAP staining. Red and magenta arrows indicate the signal for neurons and astrocytes, respectively. (**F** and **G**) Magnification of the DG (**F**) and cerebellum (**G**) showing the localization of SEL1L and GFAP. White arrows indicate SEL1L-positive signal in granule cells (**F**) and Purkinje cells (PC) (**G**). (**H** and **I**) Representative zoomed-in images for DG (**H**) and cerebellum (**I**) showing separated channels for SEL1L, NeuN, and GFAP staining. The molecular layer (ML) and granular layer (GL) are indicated as reference points in panel (**I**). Arrows, neurons (red) and astrocytes (magenta). (**J**-**M**) Quantitation of SEL1L signal intensity in neurons (red) and astrocytes (magenta) in cortex (**J**), CA1 (**K**), DG (**L**), and cerebellum (M) regions of 5-week-old mice (total of 180-200 cells from *n*=3 mice each cohort). Data are shown as the mean ± SEM. \**P*< 0.05, \*\**P*< 0.01, and \*\*\**P*< 0.001 by *t*-test (**J**-**M**). Scale bar: 2 mm (**A**); 100 μm (**B**-**C**, and **F**); 50 μm (**G**); 20 μm (**D**-**E**, and **H**-**I**).

### Generation and validation of a neuron-specific *Sel1L-*deficient mouse model

To investigate the role of the SEL1L-HRD1 ERAD pathway in neurons, we generated *Sel1L^SynCre^* mice by crossing *Sel1L*-floxed (*Sel1L^f/f^*) animals with a Synapsin I promoter-driven Cre line (40). The Synapsin I promoter becomes active around embryonic day 14-16, depending on the brain region, and remains active in postmitotic neurons (40). To assess SEL1L deletion in the brain, we performed confocal microscopy following immunofluorescence staining for SEL1L and the ER marker KDEL on coronal sections from 5-week-old mice (Figure 2). KDEL signal is known to increase upon ERAD deficiency as part of a compensatory response (41). In WT *Sel1L^f/f^* littermates, SEL1L co-localized with KDEL throughout the cortex and hippocampus (yellow, Figure 2A-B, upper panels, and F, left panel). In contrast, *Sel1L^SynCre^* mice showed loss of SEL1L in subsets of neurons, accompanied by a marked increase in KDEL staining in both regions (red, Figure 2A-B, F). These regions are critical for motor function, learning, and memory (42, 43). Quantification revealed that ∼37% of cortical neurons lacked detectable SEL1L (Figure 2C), while deletion efficiency approached 100% in the CA3 and DG subregions of the hippocampus (arrows, Figure 2D, F, G). Interestingly, SEL1L expression in the CA1 region was largely preserved (Figure 2E-F). Western blot analysis confirmed reduced SEL1L levels in both cortex and hippocampus (Supplemental Figure 2). HRD1 protein was also decreased in the hippocampus but remained unchanged in the cortex. In contrast, OS9, a lectin and known ERAD substrate, was significantly upregulated in both regions (Figure S2A-D), indicating ERAD dysfunction. Together, these results validate the successful generation of a neuron-specific *Sel1L* knockout model, with regionally distinct deletion efficiencies and evidence of ERAD disruption in affected neurons.

**Figure 2.**
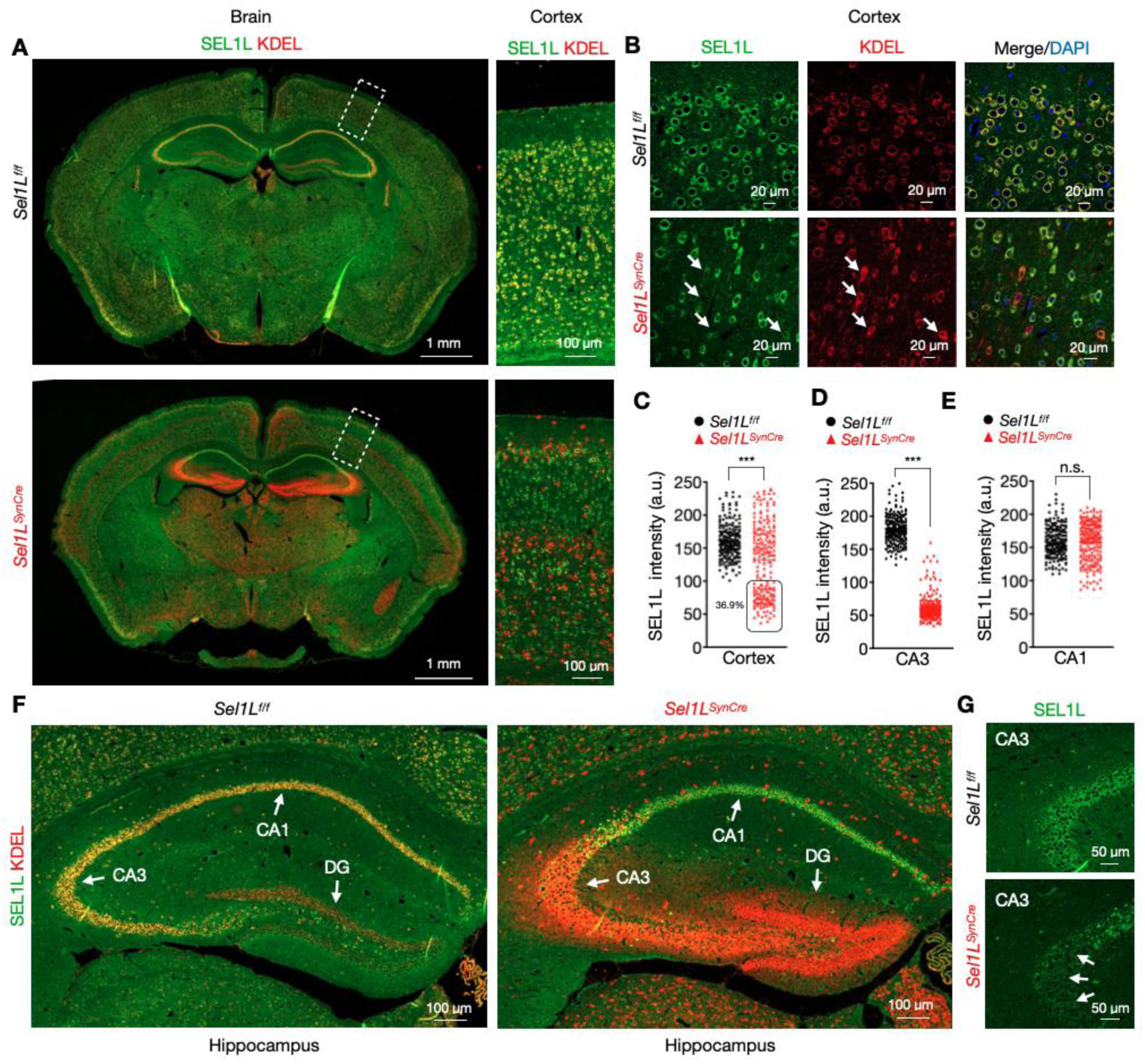
Generation of a neuronal *Sel1L*-deficient mouse model. (**A**) Representative confocal images of SEL1L (green) and KDEL (red) staining in coronal brain sections of 5-week-old wild-type (*Sel1L*^f/f^, upper panel) and *Sel1L*-deficient mice (*Sel1L*^SynCre^). The magnification of the selected region for cerebral cortex is showed in the lateral panel for each image. (**B**) Representative zoom-in images for the external layer of the cortex showing separated channels for SEL1L and KDEL staining. White arrows indicate cells with reduced levels of Sel1L. Elevated KDEL signal is used to identify *Sel1L*-deficient neurons. (**C**-**E**) Quantitation of SEL1L signal intensity in cortex (**C**), CA3 (**D**), and CA1(**E**) regions of 5-week-old mice (total of 180-200 cells from n=3 mice each cohort). (**F**) Magnification of the hippocampus for wild-type (left panel) and *Sel1L*-deficient mice (right panel). White arrows indicate the CA1, CA3, and DG regions of the hippocampus. (**G**) High-magnification images showing the reduction of SEL1L signal in the CA3 region of the hippocampus in (white arrows). Data are shown as the mean ± SEM. \**P*< 0.05, \*\**P*< 0.01, and \*\*\**P*< 0.001 by *t*-test (**C**-**E**). Scale bar: 1 mm (**A**); 100 μm (**A** lateral panel, **F**); 50 μm (**G**); 20 μm (**B**).

### Neuronal deficiency of SEL1L, but not IRE1α, leads to growth retardation and premature lethality

*Sel1L^SynCre^* mice were born at Mendelian ratios and were phenotypically indistinguishable from their littermate controls at birth. However, by two weeks of age, both male and female *Sel1L^SynCre^* mice exhibited pronounced growth retardation, with body weights reaching only ∼50% of those of WT littermates by two months of age (Figure 3, A-C). These mice also demonstrated increased post-weaning mortality, with a median survival of 9 weeks (Figure 3, D-F). In terminal stages, *Sel1L^SynCre^* mice developed tremors and typically died before 12 weeks of age (Video 1). Histopathological analysis of peripheral tissues – including brown and white adipose tissue, liver, skeletal muscle, kidney, and pancreas – revealed no overt abnormalities (Supplemental Figure 3). To determine whether disruption of the unfolded protein response (UPR) elicits similar phenotypes, we generated *Ire1a^SynCre^*mice, in which IRE1α, a major UPR sensor (44), was conditionally deleted in neurons using the same Synapsin I promoter-driven Cre line. Western blotting confirmed efficient deletion of IRE1α in both cortex and hippocampus (Supplemental Figure 4A). In contrast to *Sel1L^SynCre^* mice, *Ire1a^SynCre^*mice developed normally and were indistinguishable from their WT littermates with respect to body weight, appearance, and overall health (Figure 3, G-I). Moreover, BiP expression, PERK levels, and eIF2α phosphorylation remained unaltered, indicating preserved ER homeostasis (Supplemental Figure 4, A and B). These findings establish that the SEL1L–HRD1 ER-associated degradation (ERAD) pathway, but not neuronal IRE1α signaling, is essential for maintaining neuronal homeostasis and is critically required for postnatal growth and survival.

**Figure 3.**
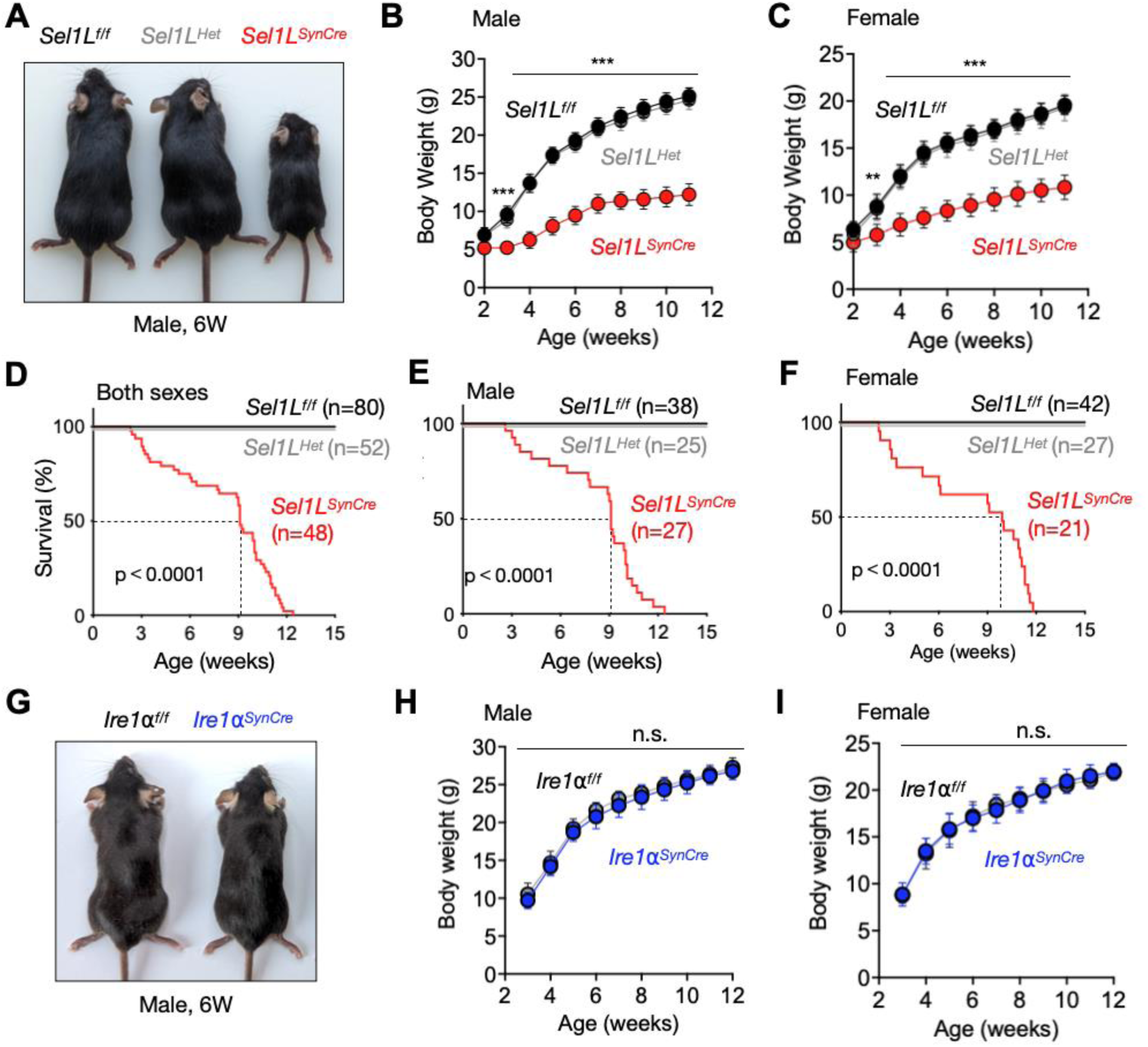
Growth retardation and premature lethality of *Sel1L*^SynCre^ mice, but not *Ire1a*^SynCre^ mice. (**A**) Representative images of 6-week-old wild-type (*Sel1L*^f/f^), heterozygote (*Sel1L*^Het^), and neuronal Sel1L-deficient (*Sel1L*^SynCre^) male mice. (**B** and **C**) Growth curve for male (**B**) and female (**C**) *Sel1L*^f/f^, *Sel1L*^Het^, and Sel1L^SynCre^ mice (2-11 weeks, *n* = 12-16 mice per group). (**D**-**F**) Kaplan-Meier mouse survival curves of both sexes (**D**), male (**E**), and female (**F**) cohorts. Overall survival was followed up for 15 weeks. The number of animals is indicated for each genotype. P value was calculated using the Log-rank test. (**G**) Representative images of 6-week-old wild-type (*Ire1⍺*^f/f^) and neuronal IRE1⍺-deficient (*Ire1⍺*^SynCre^) male mice. (**H** and **I**) Growth curve for male (**H**) and female (**I**) *Ire1⍺*^f/f^ and *Ire1⍺*^SynCre^ mice (3-12 weeks, *n* = 12-14 mice per group).

### Impaired motor activity and behavioral abnormalities in *Sel1L^SynCre^* mice

*Sel1L^SynCre^* mice began to exhibit neurological abnormalities as early as two weeks of age (Video 2). A prominent early phenotype was abnormal limb-clasping reflexes when suspended by the tail, in contrast to WT littermates, which showed normal limb extension (Figure 4, A and B). In comparison, *IRE1⍺^SynCre^* mice exhibited normal limb-clasping responses, even at 12 weeks of age (Figure 4, C and D). In the open field test, *Sel1L^SynCre^* mice demonstrated significantly reduced spontaneous locomotor activity (Figure 4E). Quantitative analysis over a 30-minute period revealed a 60% reduction in total distance traveled (Figure 4F) and a 63% decrease in time spent in the center of the arena, indicating increased anxiety-like behavior (Figure 4G). Additionally, these mice exhibited a 68% reduction in rearing frequency, an exploratory behavior in rodents (45), further supporting behavioral impairment (Figure 4H). Motor coordination was also compromised in *Sel1L^SynCre^* mice, as evidenced by an 80% reduction in latency to fall during rotarod testing (Figure 4I). In contrast, *IRE1⍺^SynCre^* mice displayed no significant deficits in locomotor activity or behavior (Figure 4, J-L). Collectively, these data indicate that SEL1L expression in neurons is essential for maintaining normal motor coordination and behavioral responses, including anxiety, in mice.

**Figure 4.**
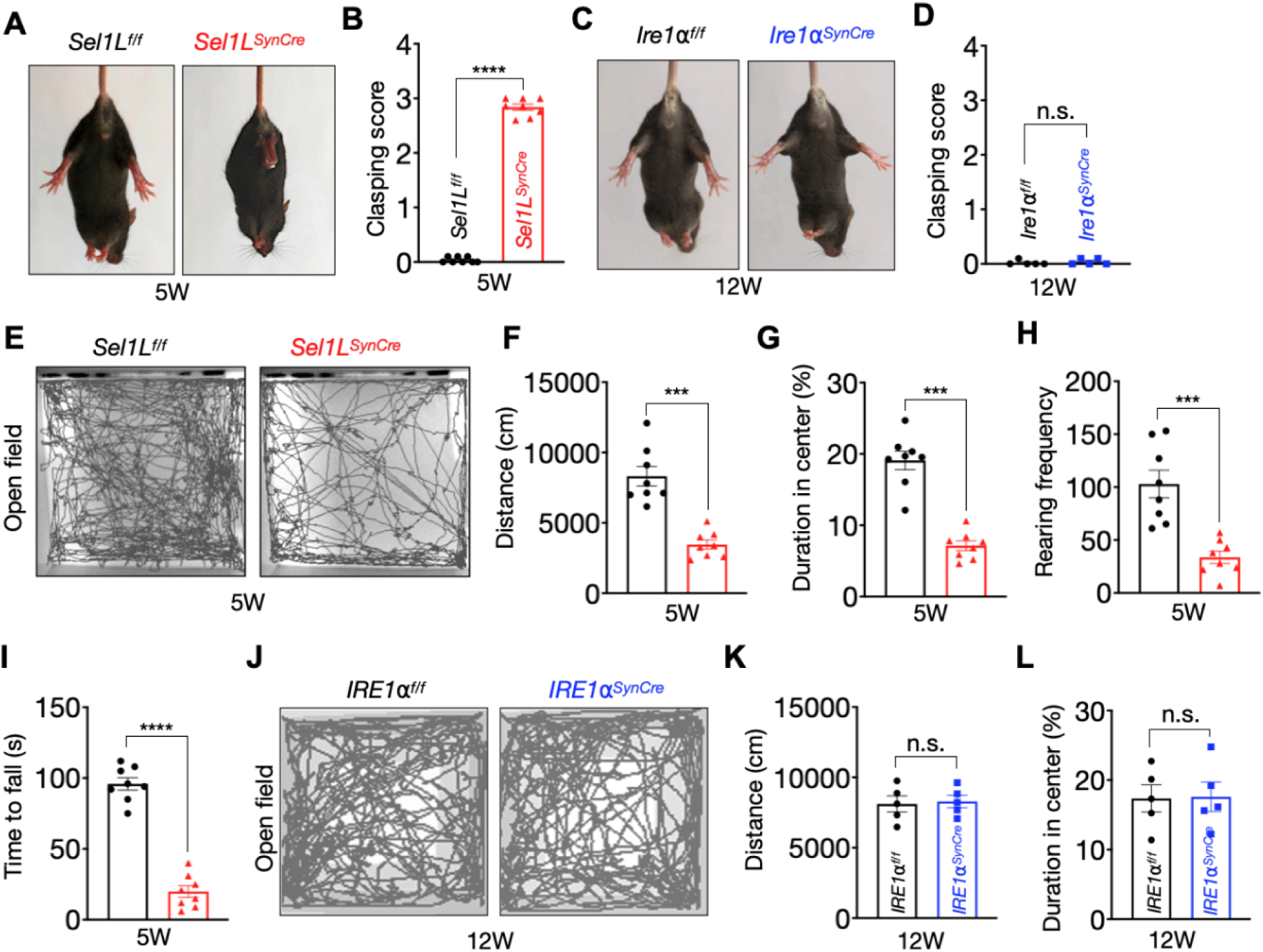
Impaired motor activity and behavior alterations of *Sel1L*^SynCre^ mice, but not *Ire1a*^SynCre^ mice. (**A**) Representative images of 5-week-old wild-type (*Sel1L*^f/f^) and neuronal Sel1L-deficient (*Sel1L*^SynCre^) male mice showing hindlimb-clasping reflex. Clasping score quantitation is showed in panel **B**, (5 weeks, *n* = 8 mice per group). (**C**) Representative images of 12-week-old wild-type (*Ire1⍺*^f/f^) and neuronal IRE1⍺-deficient (*Ire1⍺*^SynCre^) male mice. Clasping score quantitation is showed in panel **D** (12 weeks, *n* = 5 mice per group). (**E**) Open-field test showing representative locomotor activity traces for *Sel1L*^f/f^ and *Sel1L*^SynCre^ mice at 5 weeks of age. (**F-H**) Open field test quantification showing total distance (**F**), duration in the center (**G**), and rearing frequency. Animals were video recorded for 30 minutes and analyzed using EthoVision XT software (5 weeks, *n* = 8 for each genotype). (**I**) Motor performance on the accelerating rotarod test in *Sel1L*^f/f^ and *Sel1L*^SynCre^ mice (5 weeks, *n* = 8 mice per group). (**J**) Open-field test showing representative locomotor activity traces for *Ire1⍺*^f/f^ and *Ire1⍺*^SynCre^ mice Ire1⍺f/f at 12 weeks of age. (**K**-**L**) Open field test quantification showing total distance (**K**) and duration in the center (**L**) (12 weeks, n = 5 mice per group). The data presented are shown as mean ± SEM and were analyzed using a Student’s t test. Significance levels are indicated as follows: *p < 0.05; **p < 0.01; ***p < 0.001; ns, not significant.

### Neuronal SEL1L deficiency does not induce significant cell death

To investigate the impact of SEL1L deficiency in neurons, we compared brain size between *Sel1L^SynCre^* mice and WT littermates. Although *Sel1L^SynCre^* mice exhibited a 22% reduction in absolute brain weight compared to WT littermates, brain weight normalized to body weight was comparable between groups (Figure 5, A and B). Histological analysis using H&E staining revealed an approximately 20% reduction in cortical thickness and DG size (granular cell layer) in *Sel1L^SynCre^* mice, consistent with reduced brain size (Figure 5, C-F). No significant changes were observed in the CA1 region, in line with the lack of SEL1L deletion there (Supplemental Figure 5, A and B). Despite these structural alterations, neuronal density assessed by immunofluorescence staining for the neuronal marker NeuN was slightly increased in both the cortex and DG of *Sel1L^SynCre^* mice (Figure 5, G-J). TUNEL assays showed no significant differences in DNA fragmentation between *Sel1L^SynCre^* and WT mice in either region (Figure 5, K and L; Supplemental Figure 5, C and D). Furthermore, levels of cleaved caspase-3, a marker of apoptosis, were unchanged in the cortex and hippocampus (Supplemental Figure 5E and F). Together, these findings indicate that neuronal SEL1L deficiency does not reduce neuronal density or induce apoptosis at 9 weeks of age.

**Figure 5.**
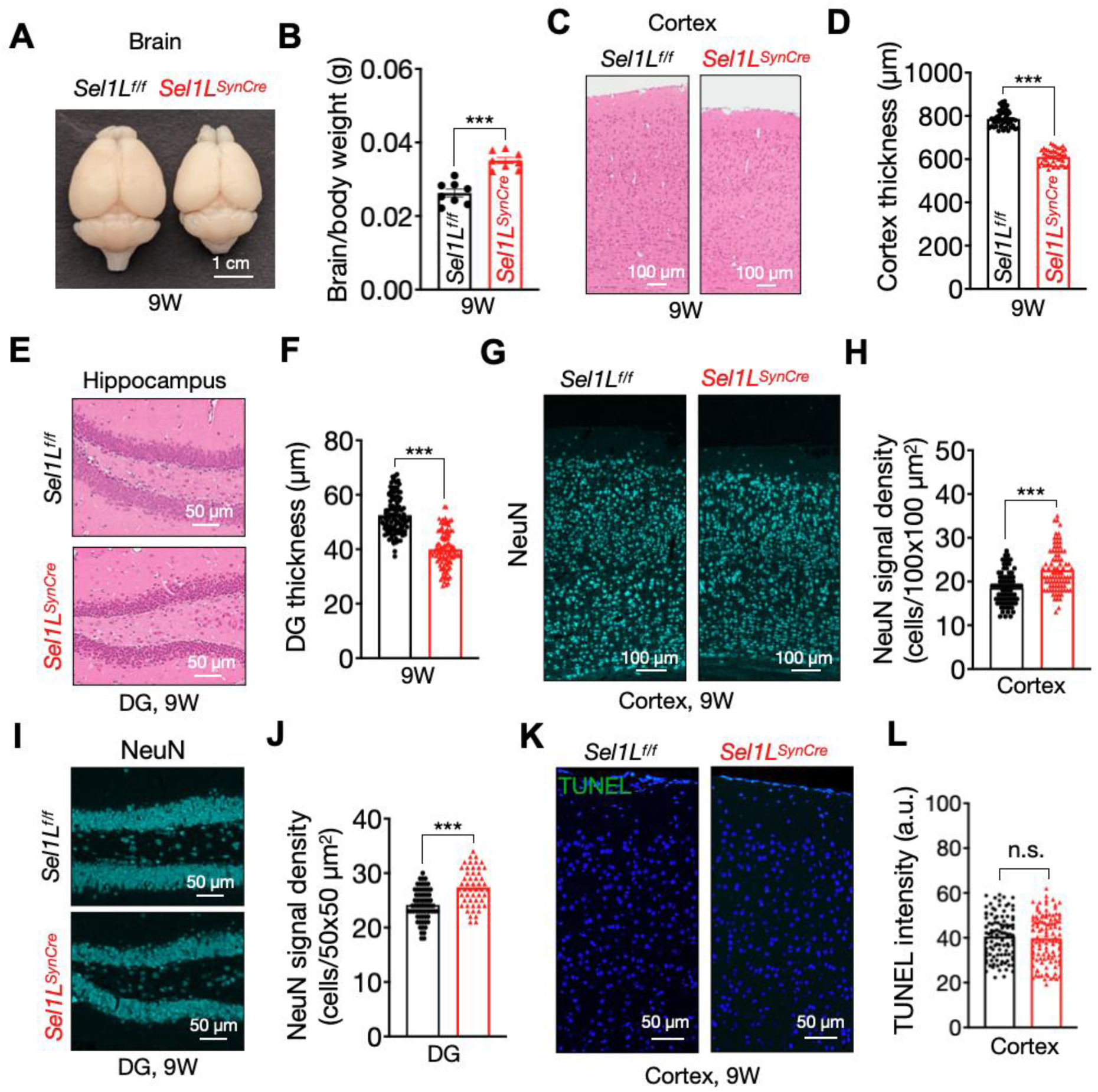
Neuronal SEL1L deficiency does not induce significant cell death. (**A**) Representative images of brains from *Sel1L*^f/f^ and *Sel1L*^SynCre^ mice at 9 weeks of age. (**B**) Whole brain weight for both genotypes normalized by body weight (n = 8 mice per group). (**C**) Hematoxylin and eosin staining of the cerebral cortex from *Sel1L*^f/f^ and *Sel1L*^SynCre^ mice at 9 weeks of age. Quantitation of cortical thickness is shown in panel **D** (50-60 measurements; n = 3 mice per group). (**E**) Hematoxylin and eosin staining of the DG region in the hippocampus at 9 weeks of age. Quantitation of the granular layer thickness in the DG is shown in panel **F** (50-60 measurements; n = 3 mice per group). (**G**) Representative confocal images of NeuN staining in cerebral cortex sections from *Sel1L*^f/f^ and *Sel1L*^SynCre^ mice at 5 weeks of age. Quantitation of NeuN-positive cells (neurons) density in the cortex is shown in panel **H** (50-60 measurements; n = 3 mice per group). (**I**) Representative confocal images of NeuN staining in the DG region of the hippocampus at 5 weeks of age. Quantitation of NeuN-positive cells (neurons) density in the granular layer is shown in panel **J** (50-60 measurements; n = 3 mice per group). (**K**) Analysis of DNA fragmentation by TUNEL assay in cerebral cortex sections from *Sel1L*^f/f^ and *Sel1L*^SynCre^ mice at 5 weeks of age. Quantification of the average signal within a 100 x 100 µm^2^ tissue area is shown in panel **L** (100-120 measurements; n = 3 mice per group). The data presented are shown as mean ± SEM and were analyzed using a Student’s t test. Significance levels are indicated as follows: *p < 0.05; **p < 0.01; ***p < 0.001; ns, not significant.

### Mild activation of the UPR in *Sel1L^SynCre^* mice

To assess ER homeostasis in *Sel1L^SynCre^* mice, we evaluated the expression and activity of key UPR sensors and their downstream effectors, as previously described (46, 47). Western blot analysis revealed a marked increase in IRE1α protein levels in *Sel1L^SynCre^*cortical lysates compared with WT controls (Figure 6, A and B), consistent with prior findings that IRE1α is a substrate of SEL1L-HRD1 ERAD (48). IRE1α-mediated splicing of *Xbp1* mRNA was elevated in *Sel1L^SynCre^* mice, increasing from ∼ 1% in WT to ∼ 7% in mutant mice (Figure 6, C and D), indicative of modest activation of the IRE1α arm of the UPR. Activation of the PERK pathway was also modest, as evidenced by a modest increase in PERK and eIF2α phosphorylation (Figure 6, E and F). Transmission electron microscopy (TEM) revealed a ∼ 3-fold increase in ER lumen width in neurons from *Sel1L^SynCre^*mice, indicative of ER dilation (Figure 6, G and H; Supplemental Figure 6). These findings indicate mild UPR activation, accompanied by ultrastructural changes in the ER.

**Figure 6.**
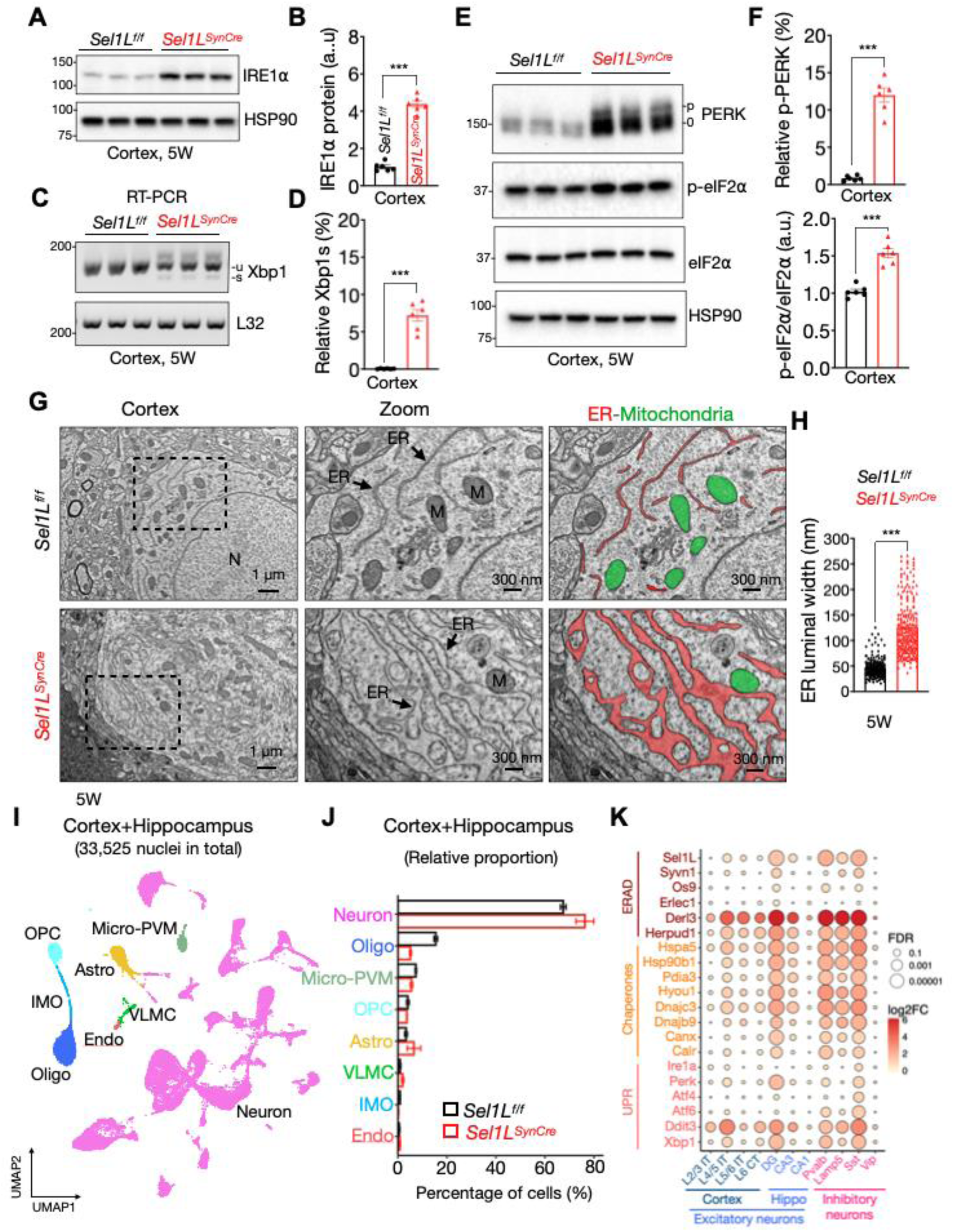
Partial UPR activation in Sel1L-deficient neurons. (**A**) Western blot analysis of IRE1⍺ in cortical tissues of *Sel1L*^f/f^ and *Sel1L*^SynCre^ mice at 5 weeks of age, with quantitation shown in **B** (*n* = 6 mice per group). Values shown are in kDa. (**C**) RT-PCR of *Xbp1* mRNA splicing from cortical tissues of 5-week-old mice. u and s indicate the unspliced and spliced form of *Xbp1*. Quantitation of the spliced form of Xbp1 (Xbp1s) is shown in **D** (n = 6 mice per group). (**E**) Western blot analysis of the PERK sensor, including the eIF2⍺ pathway in cortical tissues of mice at 5 weeks of age, with quantitation shown in **F** (*n* = 6 mice per group). (**G**) Representative TEM images from *Sel1L*^f/f^ and *Sel1L*^SynCre^ mice at 5 weeks of age. Endoplasmic reticulum (ER) and mitochondria (M) are highlighted in red and green colors respectively. Quantitation of luminal ER width is shown in **H** (200-250 measurements from 12 cells, *n* = 3 mice per group). (**I**) UMAP plot showing cell types identified by unsupervised clustering of single-nucleus RNA-seq data from combined cortex and hippocampus tissue of *Sel1L*^f/f^ and *Sel1L*^SynCre^ mice at 5 weeks of age. Each dot represents a single nucleus. (**J**) Relative abundance of cells across different clusters in *Sel1L*^f/f^ and *Sel1L*^SynCre^ mice based on single-nucleus RNA sequencing. Cell types include neurons, oligodendrocytes (Oligo), microglia and perivascular cells (Micro-PVM), oligodendrocyte precursor cells (OPC), astrocytes (Astro), vascular and leptomeningeal cells (VLMC), inflammatory mononuclear cells (IMO), and endothelial cells (Endo). (**K**) Relative abundance of mRNAs based on single-nucleus RNA sequencing for genes associated with ER quality control pathways, including ERAD, chaperones, and the UPR, in cortical and hippocampal tissue from *Sel1L*^f/f^ and *Sel1L*^SynCre^ mice at 5 weeks of age. Clustering of excitatory and inhibitory neurons was performed using previously published cell-type specific gene markers. Data in (B, D, F, H) are presented as mean ± SEM and were analyzed using a Student’s *t* test. Data in (K) was analyzed using a two-way ANOVA with FDR correction. Data is shown as log2 fold change (FC). Significance levels are indicated as follows: *p < 0.05; **p < 0.01; ***p < 0.001.

### Single-nucleus RNA sequencing reveals cellular adaptation in *Sel1L^SynCre^* neurons

To further investigate the impact of Sel1L deletion at single-cell resolution, we performed single-nucleus RNA sequencing (snRNA-seq) on cortical and hippocampal tissues from *Sel1L^SynCre^* and WT mice. A total of 33,525 nuclei were analyzed and classified into major brain cell types, including neurons, oligodendrocytes, microglia/perivascular macrophages (micro-PVM), oligodendrocyte precursor cells (OPCs), and astrocytes, based on canonical marker genes (Figure 6I and Supplemental Figure 7, A and B) (39). While the overall distribution of cell types was similar between genotypes, *Sel1L^SynCre^* mice exhibited a modest increase in neuronal representation and a reduction in oligodendrocyte proportions, with astrocyte and microglial populations remaining unchanged (Figure 6J).

We next subclustered neuronal nuclei from cortex and hippocampus and annotated their regional and cellular identities using well-established transcriptomic markers (Supplemental Figure 7, C and D) (39). Among inhibitory neurons, GABAergic subtypes expressing *Pvalb*, *Sst*, and *Lamp5* exhibited the most pronounced transcriptional alterations in *Sel1L^SynCre^*mice (Supplemental Figure 8, A and B). Within excitatory populations, deep-layer cortical projection neurons (L4/5 IT CTX) and dentate gyrus (DG) granule cells were among the most affected subtypes (Supplemental Figure 8, A and B). Within these vulnerable neuronal clusters, we assessed expression of genes involved in ER protein quality control (Figure 6K). Several ERAD genes, including *Derl3* (Derlin-3) and *Herpud1* were upregulated, along with chaperones such as *Hspa5* (BiP), *Hsp90b1* (GRP94), and *Hyou1* (GRP170), particularly in DG granule cells and *Pvalb+* and *Sst+* inhibitory neurons, where the response to ERAD deficiency appears more pronounced (Figure 6K). Additional ER chaperones, including *Canx* (calnexin), *Calr* (calreticulin), and *Pdia3* (ERp57), were also upregulated in the same neuronal populations (Figure 6K). Moreover, select UPR components, such as *Eif2ak3* (Perk), *Xbp1*, and *Ddit3* (CHOP), were elevated, whereas others, including *Ern1* (*Ire1a*), *Atf6*, and *Atf4* were unchanged (Figure 6K). Together, these data uncover a cell type-specific transcriptional response to ERAD deficiency in *Sel1L^SynCre^*mice, marked by upregulation of ER protein quality control pathways in distinct excitatory and inhibitory neuronal populations.

### Neuronal *Sel1L* deficiency activates one-carbon metabolism

To identify pathways dysregulated by neuronal loss of Sel1L, we performed KEGG pathway enrichment analysis using our snRNA-seq dataset from cortical and hippocampal regions. In addition to expected upregulation of pathways related to protein synthesis and ER quality control, we observed significant enrichment of one-carbon metabolism and amino acid biosynthesis pathways, including “biosynthesis of amino acids”, “glycine, serine, and threonine metabolism”, and “one carbon pool by folate” (highlighted in red, Figure 7A). One-carbon metabolism encompasses the folate cycle, methionine cycle, and transsulfuration pathway (Figure 7B), which collectively support methylation reactions, nucleotide synthesis, and redox homeostasis (35).

**Figure 7.**
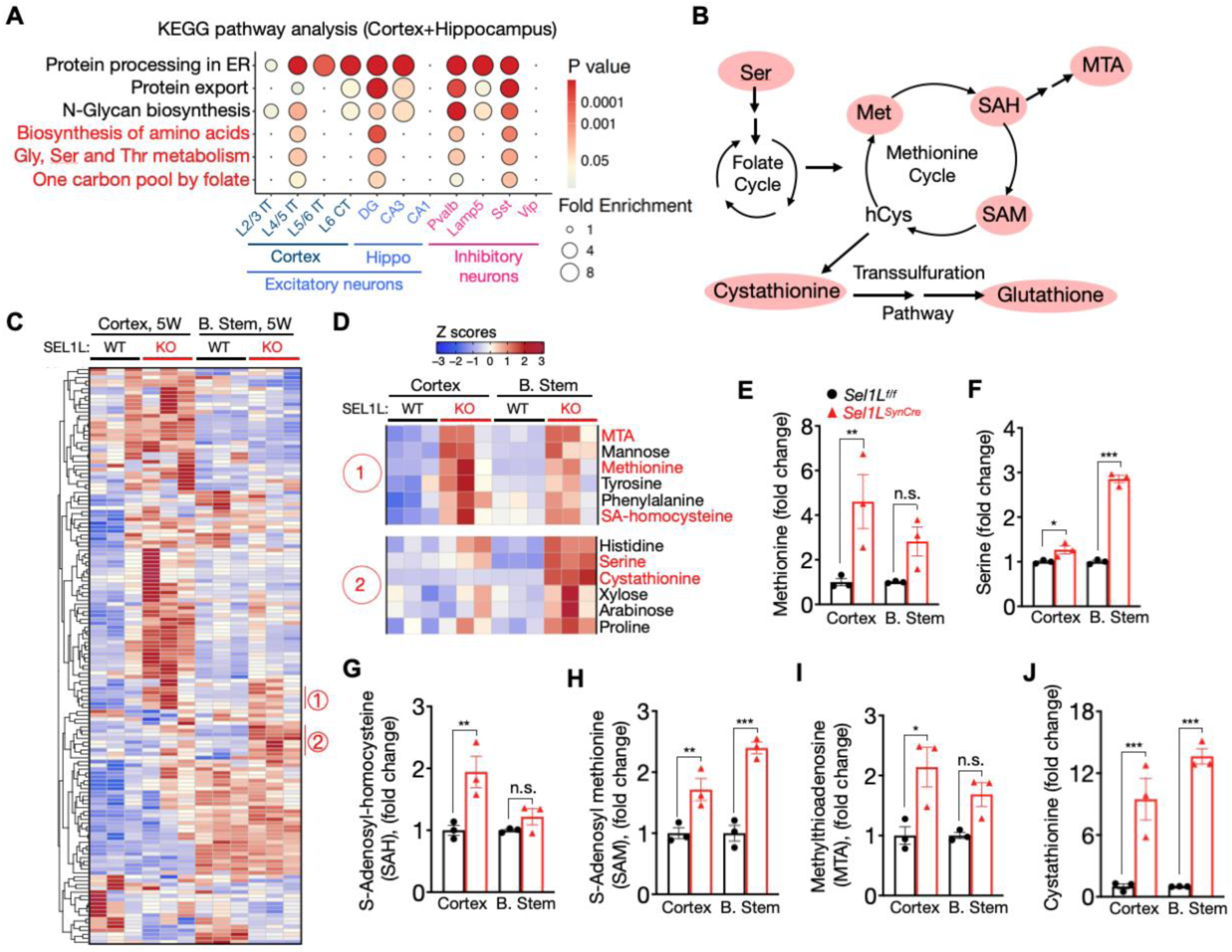
Upregulation of one-carbon metabolism pathways in the brain of *Sel1L*^SynCre^ mice. (**A**) KEGG pathway enrichment analysis of differentially expressed genes in *Sel1L*^SynCre^ mice, separated by neuronal subtypes isolated from the cortex and hippocampus at 5 weeks of age. (**B**) Diagram illustrating one-carbon metabolism, including folate, methionine, and transsulfuration pathways. Key metabolites analyzed include methionine (Met), serine, S-adenosyl-homocysteine (SAH), A-adenosyl methionine (SAM), methylthioadenosine (MAT), cystathionine, and glutathione. (**C**-**J**) Metabolomics analysis of cortex and brain stem tissues isolated from *Sel1L*^f/f^ (WT) and Sel1L^SynCre^ (KO) mice at 5 weeks of age. Panel **D** highlights selected regions showing changes in metabolites related to one-carbon metabolism. (**E**-**J**) Fold-change plots showing individual metabolites from the cortex and brain stem of *Sel1L*^f/f^ and *Sel1L*^SynCre^ mice (*n* = 3 animals per brain region). The data presented are shown as mean ± SEM and were analyzed using a two-way ANOVA with FDR correction. Significance levels are indicated as follows: *p < 0.05; **p < 0.01; ***p < 0.001.

To validate these findings, we performed metabolomic profiling in two anatomically distinct brain regions: the cortex and the brain stem (pons and medulla). Deletion of SEL1L in the brain stem was confirmed by Western blot (Supplemental Figure 9, A and B). Notably, heatmap analysis revealed a consistent increase in metabolites associated with one-carbon metabolism in both regions of *Sel1L^SynCre^* mice compared with WT controls (Figure 7, C and D). Specifically, methionine, serine, S-adenosylhomocysteine (SAH), S-adenosylmethionine (SAM), methylthioadenosine (MTA), and cystathionine were significantly increased (Figure 7, E-J). In contrast, levels of both reduced and oxidized glutathione remained unchanged in the cortex and brainstem (Supplemental Figure 9, C and D). These metabolite changes pointed to possible potential changes in methylation potential, redox balance, and epigenetic regulation, key features associated with neurometabolic and neurodevelopmental disorders (49). Together, these results indicate that neuronal *Sel1L* deficiency induces broad activation of one-carbon metabolism.

### Activation of ATF4 and one-carbon metabolic genes in *Sel1L^SynCre^* neurons

The integrated stress response (ISR), particularly the eIF2α-ATF4 axis, plays a central role in adapting to cellular stress by modulating amino acid metabolism and antioxidant responses (28, 50). Consistent with the ISR activation, we observed a ∼40% increase in eIF2α phosphorylation (Figure 6, E and F), along with 7-fold and 5-fold elevation in ATF4 protein levels in the cortex and hippocampus of *Sel1L^SynCre^*mice, respectively (Figure 8, A and B). As ATF4 is also a known transcription regulator of enzymes involved in one-carbon metabolism (51, 52), we next analyzed the expression of one-carbon metabolism genes in the snRNA-seq dataset (Figure 8C). Mitochondrial genes including *Mthfd2* (methylenetetrahydrofolate dehydrogenase 2) and *Shmt2* (serine hydroxymethyltransferase 2) were significantly upregulated, along with moderate increases in *Adh1l2* (alcohol dehydrogenase 1-like protein 2) and *Mthfd1l* (methylenetetrahydrofolate dehydrogenase 1-like) (Figure 8C). Genes in the phosphoserine pathway, *Phgdh* (phosphoglycerate dehydrogenase), *Psph* (phosphoserine phosphatase), and *Psat1* (phosphoserine aminotransferase 1), were also elevated, consistent with increased serine biosynthesis from glucose (53). In contrast, expression of genes involved in cytosolic folate and methionine metabolism—including *Mthfr* (methylenetetrahydrofolate reductase), *Shmt1* (serine hydroxymethyltransferase 1), *Mthfd1* (methylenetetrahydrofolate dehydrogenase 1), *Mat2a* (methionine adenosyltransferase 2A), *Ahcy* (adenosylhomocysteinase), and *Mtr* (methionine synthase)—as well as *G6pdx* (glucose-6-phosphate dehydrogenase X-linked), a key enzyme in the pentose phosphate pathway (54), remained unchanged (Figure 8C). These data support a model in which neuronal *Sel1l* deficiency activates the ATF4 pathway and selectively enhances mitochondrial folate cycle and serine biosynthesis of the one-carbon metabolism pathway (Figure 8D).

**Figure 8.**
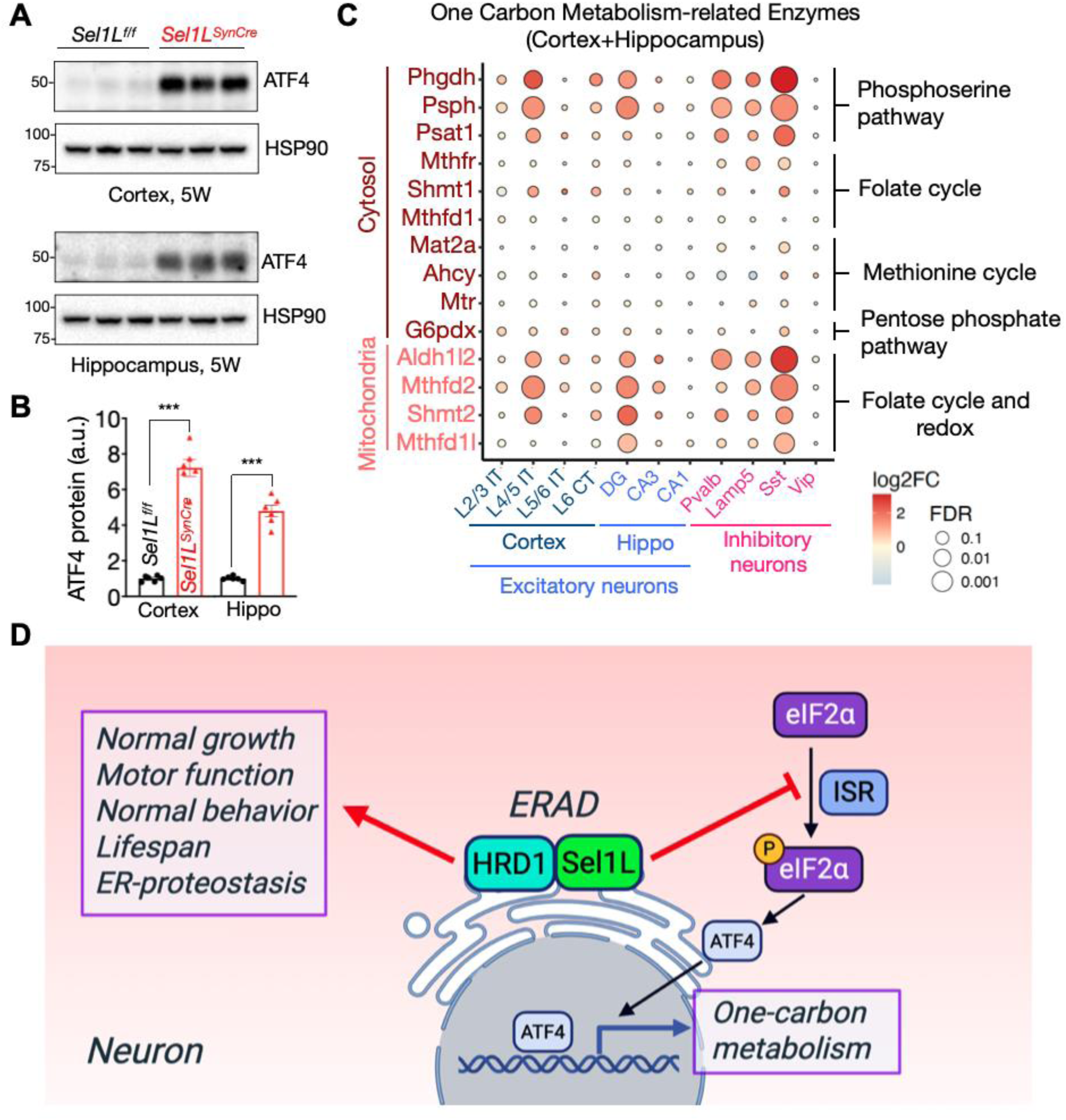
Activation of ATF4 and one-carbon metabolism enzymes in the brain of *Sel1L^SynCre^* mice. (**A**) Western blot analysis of ATF4 in cortical and hippocampal tissues from *Sel1L*^f/f^ and *Sel1L*^SynCre^ mice at 5 weeks of age, with quantitation shown in **B** (*n* = 6 mice per group). (**C**) Relative mRNA abundance of one-carbon metabolism genes based on single-nucleus RNA sequencing of cortical and hippocampal tissue from *Sel1L*^f/f^ and *Sel1L*^SynCre^ mice at 5 weeks of age. (**D**) Proposed model illustrating the role of ERAD in neurons. Proper ERAD function is essential for maintaining growth, motor function, behavior, and lifespan in mice by preserving ER proteostasis under basal conditions and preventing activation of the integrated stress response (IRS). Loss of ERAD leads to phosphorylation of eIF2⍺, increased ATF4 levels, and activation of one-carbon metabolism pathways. Positive and inhibitory regulatory effects are indicated by red arrow and blunt-ended line, respectively. Data in (B) are presented as mean ± SEM and differences between Sel1L^f/f^ and Sel1L^SynCre^ groups were analyzed using a Student’s *t* test. The data presented (C) was analyzed using a two-way ANOVA with FDR correction. Significance levels are indicated as follows: *p < 0.05; **p < 0.01; ***p < 0.001.

## DISCUSSION

Our findings establish that the SEL1L-HRD1 ERAD complex is essential for central nervous system function, contributing to normal growth, behavior, motor control, and survival in mice. In *Sel1L^SynCre^* mice, we observed neurological impairments and early death, accompanied by a marked activation of the one-carbon metabolism pathway—revealing an unexpected link between ERAD deficiency, neuronal dysfunction, and metabolic reprogramming. These findings, together with recent reports of SEL1L-HRD1 disease variants in ENDI patients (26, 27) and studies involving conditional deletion of *Sel1l* in adult neurons (55), underscore the essential role of this ER quality control pathway in sustaining neuronal homeostasis. Moreover, we demonstrate that SEL1L deficiency leads to only mild activation of the UPR, and that neuron-specific deletion of *Ire1a* using the same Cre driver fails to recapitulate the motor and behavioral deficits seen in *Sel1L^SynCre^*mice. These results suggest that UPR activation alone is insufficient to explain the observed phenotypes and point to additional substrate-dependent mechanisms underlying neuronal vulnerability.

Motor impairment is a consistent phenotype in ERAD-deficient models (55, 56), suggesting that motor neurons and associated neural circuits are particularly dependent on intact ERAD function. While previous work demonstrated that Purkinje cell-specific deletion of Sel1L results in progressive cerebellar ataxia and neurodegeneration (34), we did not detect a marked reduction of SEL1L in cerebellar Purkinje cells in *Sel1L^SynCre^*mice, indicating relative sparing of the cerebellum in this model. To define the specific contribution of motor neurons to the observed motor deficits, targeted deletion of Sel1L in motor neurons will be necessary. In addition, snRNA-seq analysis revealed a selective reduction in mature oligodendrocytes in *Sel1L^SynCre^* mice, whereas oligodendrocyte precursor cells (OPCs) were largely preserved. Given that neuronal activity promotes OPC differentiation and myelination (57, 58), these findings raise the possibility that neuronal dysfunction may impair oligodendrocyte maturation. Future studies focusing on neuron-glia signaling, myelin ultrastructure, and circuit-level connectivity will be essential to dissect the multifactorial basis of motor impairment in the context of neuronal ERAD deficiency.

By integrating two complementary and unbiased omics approaches – single-nucleus RNA sequencing and metabolomics – we identified a prominent molecular hallmark of *Sel1L^SynCre^*mice: robust activation of one-carbon metabolism. This pathway is critical for nucleotide biosynthesis, amino acid metabolism, redox homeostasis, and epigenetic regulation (35). Notably, elevated levels of SAM and SAH suggest a disruption in methylation balance (59). As SAM is the universal methyl donor and SAH a potent inhibitor of methyltransferases, an increased SAH/SAM ratio is often associated with global DNA hypomethylation and widespread transcriptional dysregulation (60). The accumulation of upstream metabolites, including serine, methionine, and cystathionine, further supports dysregulation in redox and methylation dynamics (61), which may contribute to the neurological deficits observed in *Sel1L^SynCre^* mice. While activation of the PERK pathway may contribute to these changes, we cannot exclude the involvement of other ISR arms, such as GCN2, which responds to amino acid deprivation, or HRI, which senses mitochondrial dysfunction (62), as well as the potential role of oxidative stress (63). Whether these metabolic alterations represent compensatory responses to chronic proteostatic stress or actively drive disease pathogenesis remains unclear. Nonetheless, our findings reveal a critical role for the SEL1L–HRD1 ERAD complex in regulating neuronal metabolic homeostasis and identify one-carbon metabolism as a potential therapeutic target. This model provides a powerful platform for dissecting the molecular and metabolic basis of ERAD-associated neurodevelopmental disorders and for evaluating targeted metabolic or genetic interventions.

## MATERIALS AND METHODS

### Sex as a biological variable

Our study examined male and female animals, and similar findings are reported for both sexes.

### Mice

Neuron-specific Sel1L deficient mice (Sel1L1SynCre) were generated by breeding the Synapsin I-Cre mice on the C57BL/6J background (Jackson Laboratories #003966) with the Sel1Lflox/flox mice, also on the C57BL/6J background (1). Age-and sex-matched littermates were housed in a temperature-controlled room on a 12-h light/dark cycle.

### Genotyping

Sel1LSynCre and IRE1⍺SynCre mice were routinely genotyped using PCR of genomic DNA samples obtained from ears with the following primer pairs: Sel1Lflox/flox: F: 5’-CTGACTGAGGAAGGGTCTC-3’, R: 5’-GCTAAAAACATTACAAAGGGGCA-3’; IRE1⍺flox/flox: F: 5’-CCGAGCCATGAGAAACAAGG-3’, R: 5’-CCCTGCCAGGATGGTCATGG-3’; Cre recombinase: F: 5’-ACCTGAAGATGTTCGCGATTATCT-3’, R: 5’-ACCGTCAGTACGTGAGATATCTT-3’.

### Behavioral studies

All behavior procedures were performed by investigators blinded to the genotypes. For hindlimb clasping assessment, mice were lifted by the tail and held over a cage for 1 min to assess abnormal hindlimb clasping, and scoring was performed as previously described (2). For the open field test, mice were placed in the center of the arena, and video recording began 3 seconds after the mouse was detected. Behavior was recorded for 30 minutes. At the end of the session, mice were returned to their home cage, and the apparatus was cleaned between trials. Total distance traveled, time spent in different zones, rearing, and movement were quantified using EthoVision XT (Noldus). K2. For the rotarod test, an accelerating rotarod (Model LE8500, Panlab SL) was used. Mice were trained in one session per day over four consecutive days. The rotarod started at 4 rpm and accelerated to 40 rpm at a rate of 0.2 rpm/s. The latency to fall was recorded automatically, and performance on the fifth day was analyzed using a 3-minute cutoff.

### Western blot and antibodies

Tissues were harvested and snap-frozen in liquid nitrogen. Proteins were extracted by sonication in 1% Triton X-100 buffer (50 mM Tris-HCL at pH 7.5, 150 mM NaCl, 1% Triton X-100, 1 mM EDTA) supplemented with protease inhibitor (Sigma), 1 mM DTT (Sigma) and phosphatase inhibitor cocktail (Sigma). Lysates were incubated on ice for 20 minutes and centrifuged at 16,000 g for 10 minutes. Supernatants were collected, and protein concentration were determined using the Bio-Rad Protein Assay Dye (Bio-Rad). A total of 20-30□g of protein was denatured at 95°C for 5 minutes in 1x SDS sample buffer (50 mM Tris-HCl pH 6.8, 2% sodium dodecyl sulfate, 0.01% Bromophenol blue, 10% glycerol, and 0.3 M□- mercaptoethanol). Proteins were separated by SDS-PAGE and transfer onto PVDF membranes (Fisher Scientific). Membranes were blocked in 2% BSA in Tris-buffered saline with 0.1% tween-20 (TBST) and incubated overnight at 4°C with the following primary antibodies: anti-HSP90 (Santa Cruz, sc-7947, 1:5,000), anti-SEL1L (home-made, 1:5,000), anti-HRD1 (Proteintech, 13473-1, 1:2,000), anti-OS9 (Abcam, ab109510, 1:5,000), anti-IRE1□ (Cell Signaling, 3294, 1:2,000), anti-PERK (Cell Signaling, 3192, 1:1,000), anti-p-eIF2□ (Cell signaling, 9721, 1:1000), anti-eIF2□ (Cell signaling, 9722, 1:1000), and anti-ATF4 (Cell signaling, 11815, 1:1000). After washing with TBST, membranes were incubated with HRP-conjugated secondary antibodies (Bio-Rad, 1:5,000) for 1 hour at room temperature. Signals were developed using a ECL chemiluminescence detection system (Bio-Rad), and band intensities were quantified using the ImageJ software.

### RNA preparation and RT-PCR

Total RNA was extracted from tissues using TRI Reagent and BCP phase separation reagent according to the manufacturer’s protocol (Molecular Research Center, TR118). RT-PCR for Xbp1 mRNA splicing was performed as previously described (3). The ratio of spliced Xbp1 (Xbp1s) to total Xbp1 (Xbp1u + Xbp1s) was quantified by ImageJ software. RT-PCR primer sequences were: mXbp1 F: ACGAGGTTCCAGAGGTGGAG, R: AAGAGGCAACAGTGTCAGAG; mL32 F: GAGCAACAAGAAAACCAAGCA, R: TGCACACAAGCCATCTACTCA.

### Histology

Anesthetized mice were perfused with 20 ml of 0.9% NaCl followed by 40 ml of 4% paraformaldehyde in 0.1 M PBS (pH 7.4) for fixation. Brains were dissected and post-fixed overnight in 4% paraformaldehyde in PBS at 4°C. For hematoxylin and eosin (H&E) staining, tissues were dehydrated, embedded in paraffin, and processed at the Rogel Cancer Center Tissue and Molecular Pathology Core at the University of Michigan. Cortical and hippocampal thickness was quantified on H&E-stained coronal sections using Aperio ImageScope software.

### Immunofluorescent staining

Paraffin-embedded brain sections were deparaffinized in xylene and rehydrated through a graded ethanol series (100%, 90%, 70%), followed by rinsing in distilled water. Antigen retrieval was performed by boiling the slides in a microwave using a citric acid-based antigen unmasking solution (Vector laboratories, H-3300). Sections were then incubated in blocking solution (5% donkey serum, 0.3% Triton X-100 in PBS) for 1 hour at room temperature, followed by overnight incubation at 4°C in a humidified chamber with the following primary antibodies: anti-NeuN (EMD Millipore, ABN90, 1:500), anti-GFAP (Cell Signaling, #3670, 1:100), anti-KDEL (Novus Biologicals, #97469, 1:200), and anti-SEL1L (homemade, 1:100). The next day, after three washes with PBST (0.03% Triton X-100 in PBS), sections were incubated with the corresponding Alexa Fluor-conjugated secondary antibodies (Jackson ImmunoResearch, 1:500) for 1 hour at room temperature. Slides were then mounted with VECTASHIELD mounting medium containing DAPI (Vector Laboratories, H-1500). Images were acquired using a Nikon A1 confocal microscope at the University of Michigan Microscopy and Image Analysis Core (MIAC). Images were analyzed using ImageJ software.

### TUNEL assay

Paraffin-embedded brain sections were processed for TUNEL staining using the In-Situ Cell Death detection kit (Roche, 11684795910) according to the manufacturer’s instructions. Images were acquired using a Nikon A1 confocal microscope at the University of Michigan Microscopy and Image Analysis Core (MIAC). Quantification was performed using ImageJ software.

### Transmission electron microscopy (TEM)

Mice were anesthetized and perfused with 3% glutaraldehyde and 3% formaldehyde in 0.1 M cacodylate buffer (Electron Microscopy Sciences, 16220, 15710, 11653). Cortical brain regions were dissected and cut into small pieces, and post fixed overnight at 4°C in 3% glutaraldehyde, 3% formaldehyde in 0.1 M Sorenson’s buffer (Electron Microscopy Sciences, 11682). Tissue samples were then processed, embedded, and sectioned at the University of Michigan Microscopy Core. Sections were stained with uranyl acetate and lead citrate, and high-resolution images were acquired using a JEOL 1400-plus transmission electron microscope.

### Tissue preparation, cDNA amplification, and library construction for 10X snRNA-Seq

Mice were euthanized using isoflurane and decapitated. Brains were removed from the skull and sectioned into 1 mm sagittal slices using a brain matrix. The cortex and hippocampus from 2 Sel1Lf/f and 2 Sel1LSynCre 5-week-old mice were dissected and flash-frozen in liquid nitrogen. On the day of the experiment, frozen tissues were homogenized in lysis Buffer (EZ Prep Nuclei Kit, Sigma-Aldrich) supplemented with Protector RNase Inhibitor (Sigma-Aldrich) and filtered through a 30 µm MACS strainer (Miltenyi). Filtered samples were centrifuged at 500 x g for 5 minutes at 4°C, and the nuclear pelleted was resuspended in wash buffer (10 mM Tris Buffer, pH 8.0, 5 mM KCl, 12.5 mM MgCl2, 1% BSA with RNAse inhibitor). Nuclei were filtered again and centrifuged under the same conditions. The resulting nuclei were resuspended in wash buffer containing propidium iodide (Sigma-Aldrich), and stained nuclei were sorted using a MoFlo Astrios Cell Sorter. Sorted nuclei were centrifuged at 100 x g for 5 minutes at 4°C and resuspended in wash buffer to a final concentration of 750–1,200 nuclei/μL. Reverse transcription (RT) mix was added to target approximately 10,000 nuclei, which were then loaded onto a 10× Chromium Controller chip. The Chromium Single Cell 3′ Library and Gel Bead Kit v3, Chromium Chip B Single Cell kit, and Chromium i7 Multiplex Kit were used for RT, cDNA amplification, and library preparation according to the manufacturer’s instructions. Libraries were sequenced on an Illumina NovaSeq 6000 platform with 150 nt pair ended reads.

### snRNA-Seq data analysis

Raw sequencing reads were processed through demultiplex, mapping, and analysis by the pipeline in 10X Genomics Cell Ranger, and the output data were further analyzed including integration using the Seurat package (version 4.3.0) in the R environment (version 4.2.3). After removing doublets and cells with low quality (high mitochondrial content or low sequencing depth), 33,636 cells that expressed more than 600 genes and 23,886 genes with transcripts detected in more than 3 cells were used for further analysis. The average unique molecular identifiers (UMIs) count for each cell was 9216. Unsupervised clustering was applied at a resolution of 0.2 using the top 17 dimensions of PCA using the default pipeline in the Seurat package. Cell cluster identification was based on the prior knowledge of marker genes (64). Differential gene expression analysis was conducted for each cluster between genotypes with the pseudo-bulking method using the edgeR package (version 4.4.0). The Uniform Manifold Approximation and Projection (UMAP) plots, feature plots, and heatmaps were generated in R.

### Metabolomics

Mice were euthanized using isoflurane and decapitated. Brains were removed from the skull and sectioned into 1 mm-thick sagittal slices using a brain matrix. The cortex and hippocampus were dissected, flash frozen in liquid nitrogen, and stored at -80°C. On the day of the experiment, frozen tissue samples were homogenized in cold (-80 °C) 80% methanol. Soluble metabolites were separated from the insoluble fraction by centrifugation and dried using a SpeedVac, with volumes normalized to tissue weights. Dried metabolites were reconstituted in a 1:1 methanol:water solution for LC-MS analysis. Metabolite extracts were analyzed using an Agilent Technologies Triple Quad 6470 LC-MS/MS system, equipped with a 1290 Infinity II LC Flexible Pump (Quaternary Pump), the 1290 Infinity II Multisampler, a 1290 Infinity II Multicolumn Thermostat with 6-port valve, and a 6470 triple quadrupole mass spectrometer. Agilent MassHunter Workstation Software (LC/MS Data Acquisition for 6400 Series Triple Quadrupole MS with Version B.08.02) was used for compound optimization, calibration, and data acquisition. Extracted ion chromatograms and mass spectra were manually inspected to ensure sample quality and consistent peak integration. Pathway analysis of significantly altered metabolites was performed using MetaboAnalyst (https://www.metaboanalyst.ca). Unsupervised hierarchically clustered heatmaps were generated by using R software.

### Statistics

Statistical analyses were performed in GraphPad Prism version 8.0 (GraphPad Software). Unless indicated otherwise, values are presented as mean ± standard error of the mean (SEM). All experiments have been repeated at least two to three times and/or performed with multiple independent biological samples from which representative data are shown. All datasets passed normality and equal variance tests. Statistical differences between groups were assessed using an unpaired two-tailed Student’s t-test for comparisons between two groups, or a two-way ANOVA followed by FDR correction for multiple comparisons. A *p*-value of less than 0.05 was considered statistically significant.

### Study approval

All animal procedures were approved by the Institutional Animal Care and Use Committee of the University of Michigan Medical School (PRO00010658) and the University of Virginia (PRO0004459), in accordance with the National Institutes of Health (NIH) guidelines.

## Data and material availability

The materials and reagents used are either commercially available or available upon the request. All other data are available in the main text or Supplemental Materials.

## AUTHOR CONTRIBUTIONS

M.T. designed and performed most experiments; J.H. stablished the original SEL1L^SynCre^ colony; H.W., B.P., L.L.L, A.T. and A.G. contributed to both *in vitro* and *in vivo* experiments; Y.L and A.R. performed the snRNA-seq analysis; A.S assisted with the metabolomics analysis; M.M., Z.Z., C.L., and W.D. provided insightful discussion; L.Q. directed the study. L.Q. and M.T. wrote the manuscript; all authors commented on and approved the manuscript.

## ACKNOWLEDGEMENTS

We acknowledge to the University of Michigan Animal Phenotyping Core for the open field analyses supported by P30 grants DK020572, DK089503 and 1U2CDK135066; Microscopy and Image Analysis Core (MIAC) supported by NIH NIDDK grant P30DK020572; Sasha Meshinchi at the Michigan Medicine Microscopy Core for training and advice on the TEM sample preparation and imaging; members of the Qi and Arvan laboratories at the University of Michigan Medical School for technical assistance and insightful discussions; This work was supported by NCI F32CA260735 (A.J.S.), RF1NS122060 (Z.Z.), Alzheimer’s Association (24AARG-D-NTF-1187603), 1R01NS138119 (L.Q.), and 1R01AG089640 (L.Q., Z.Z.).

