## Supplementary figures for "Neuronal SEL1L-HRD1 ERAD regulates one-carbon metabolism and is essential for motor function and survival"

Contents:

- Supplemental Figures 1-9
- Supplemental Videos 1-2

### SUPPLEMENTAL FIGURES

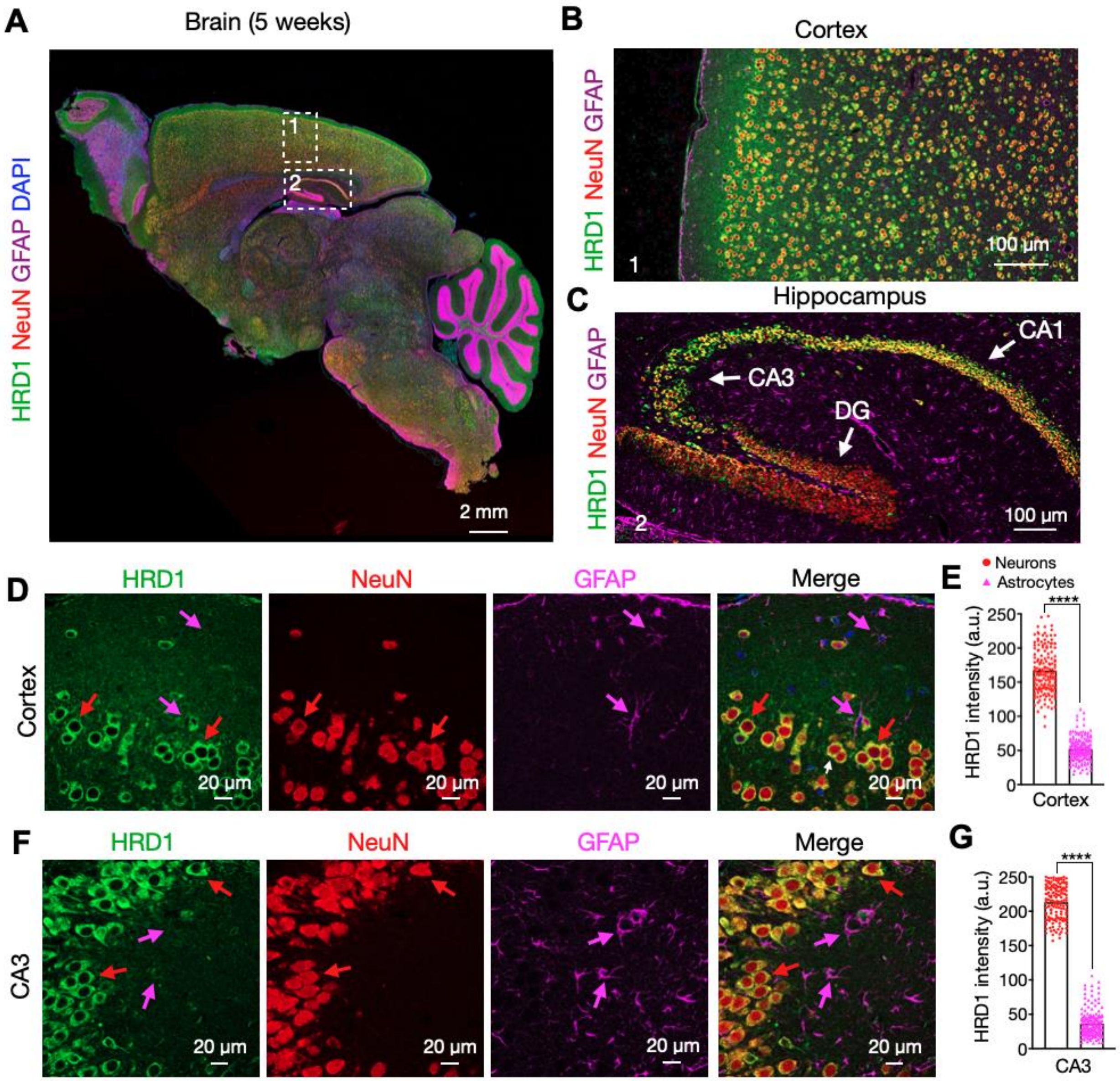


**Supplementary figure 1. HRD1 protein is highly expressed in neurons.** (**A**) Representative confocal image of HRD1 (green), NeuN (red), GFAP (magenta), and DAPI (blue) staining in a sagittal brain section of 5-week-old mouse. The magnification of the selected regions for cortex and hippocampus, are showed in panels (**B**) and (**C**), respectively. White arrows in panel (**C**) indicate the CA1, CA3, and DG regions of the hippocampus. (**D** and **F**) Representative zoomed-in images for cortex (**D**) and CA3 (**F**) regions showing separated channels for HRD1, NeuN, and GFAP staining. Red and magenta arrows indicate the signal for neurons and astrocytes, respectively. (**E** and **G**) Quantitation of HRD1 signal intensity in cortex (**E**) and CA3 (**G**) regions of 5-week-old mice (total of 180-200 cells from n=3 mice each cohort). Data are shown as the mean ± SEM. *P< 0.05, **P< 0.01, and ***P< 0.001 by t-test (**E** and **G**). Scale bar: 2 mm (A); 100 µm (B-C); 25 µm (D and **F**).

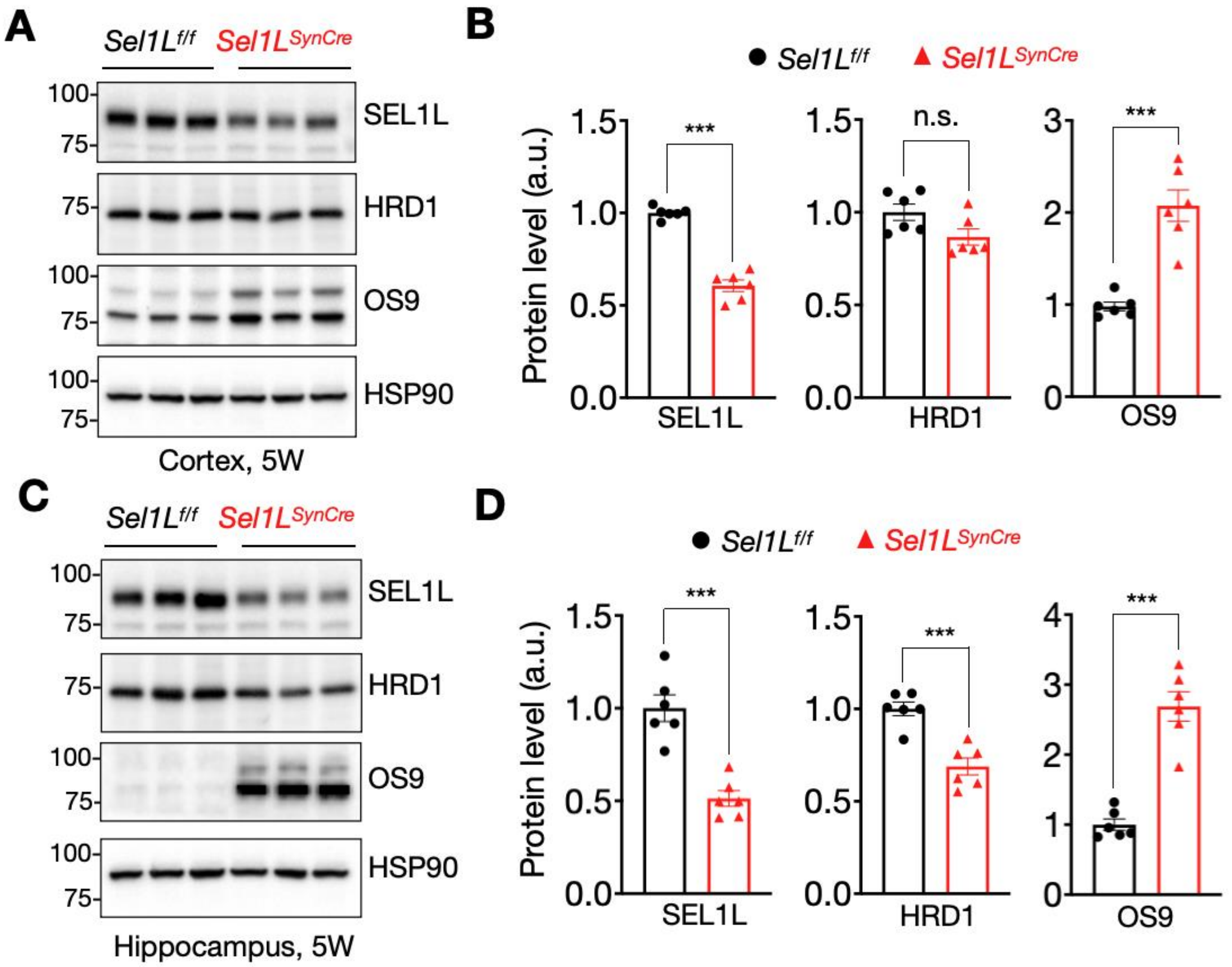


**Supplementary Figure 2. Western blot analysis of ERAD proteins in the cortex and hippocampus.** (**A**) Western blot analysis of SEL1L, HRD1, OS9 and HSP90 proteins in cortical tissues from Sel1L$^{f/f}$ and Sel1L$^{SynCre}$ mice at 5 weeks of age, with quantitation shown in (**B**) (n = 6 mice per group). (**C**) Western blot analysis of SEL1L, HRD1 and OS9 proteins in the hippocampus, with quantitation shown in (**D**) (n = 6 mice per group). Data in (B) and (D) are presented as mean ± SEM and were analyzed using an unpaired two-tailed Student's *t* test. Significance levels are indicated as follows: $*p < 0.05$; $**p < 0.01$; $***p < 0.001$.

.

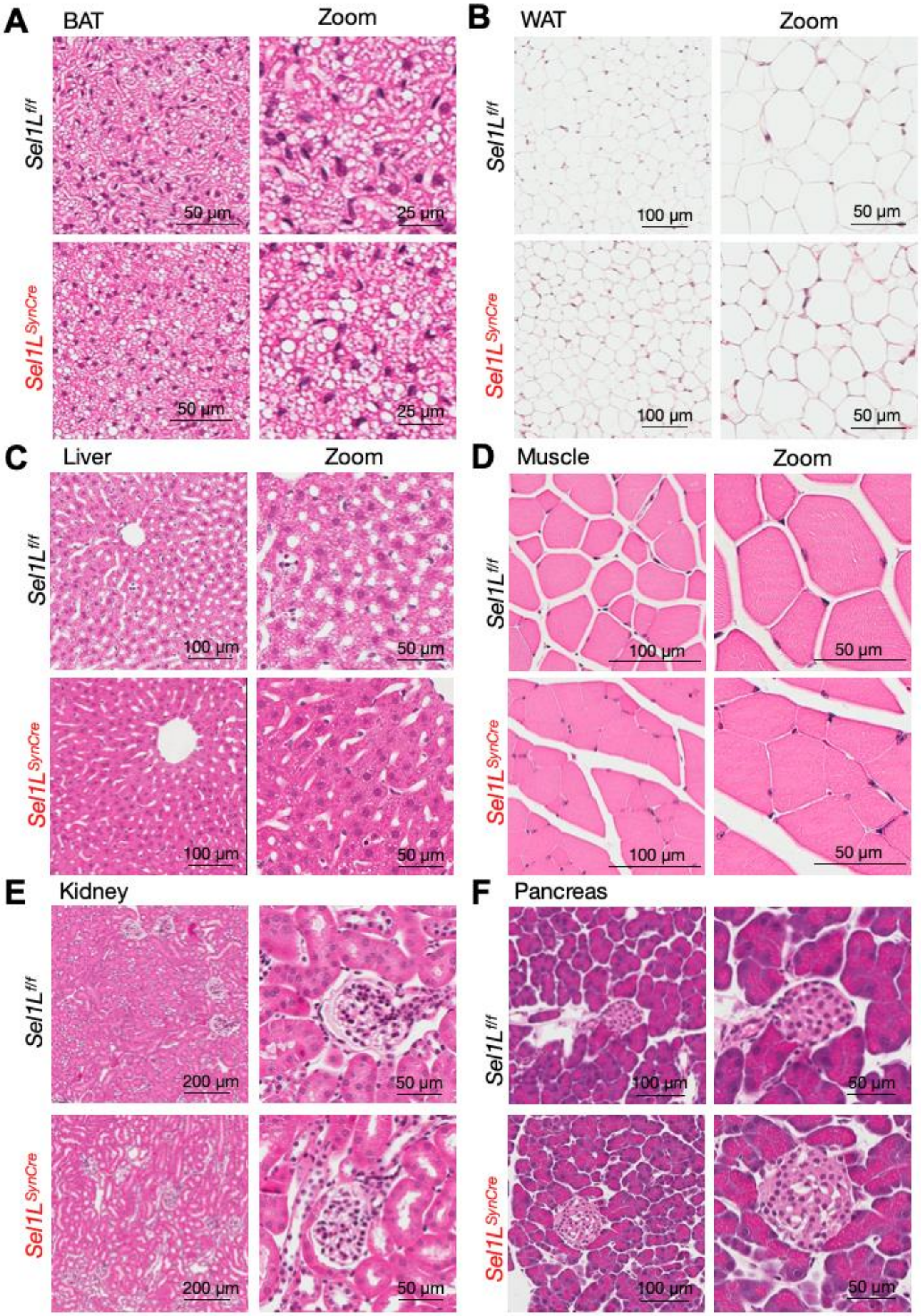


**Supplementary Figure 3. Histology of peripheral tissues in Sel1L$^{SynCre}$ mouse. (A-F)** Hematoxylin and eosin staining of brown adipose tissue (BAT) (**A**), white adipose tissue (WAT) (**B**), liver (**C**), muscle (**D**), kidney (**E**), and pancreas (**F**) from Sel1L$^{f/f}$ and Sel1L$^{SynCre}$ mice at 9 weeks of age.

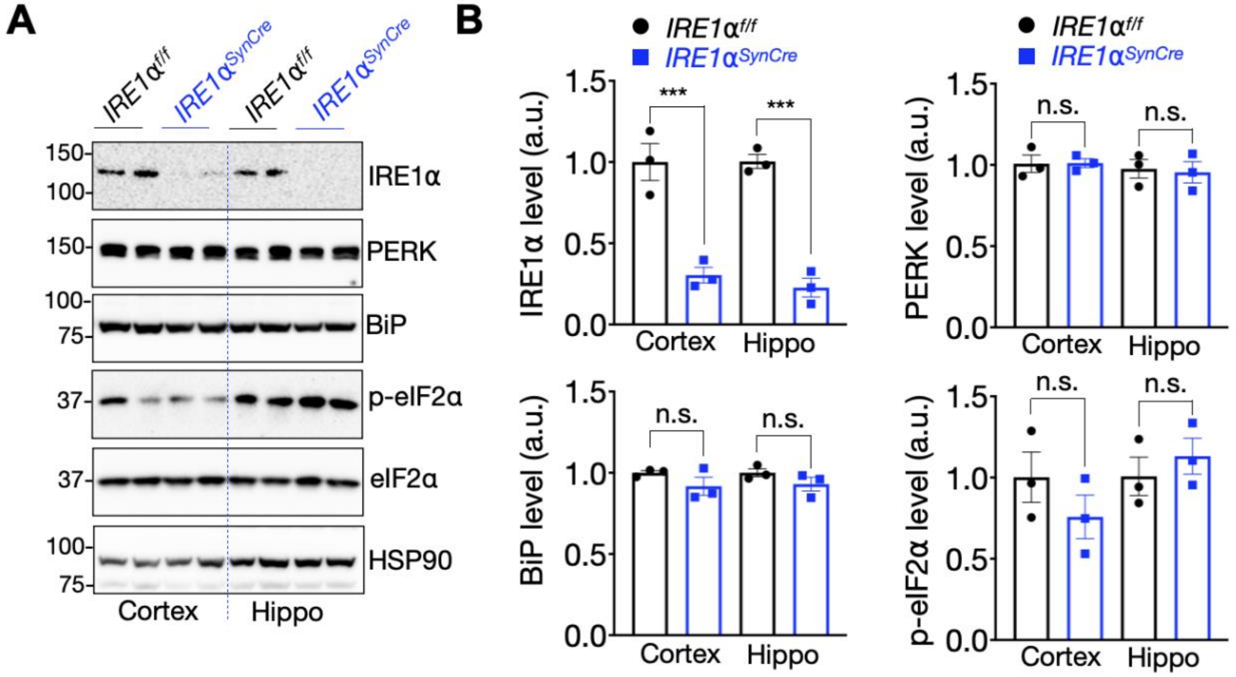


**Supplementary Figure 4. Generation of a neuronal IRE1α-deficient mouse model.** (**A**) Western blot analysis of IRE1α, PERK, Bip, and eIF2α proteins in cortical and hippocampal tissue from wild-type (IRE1$^{f/f}$) and neuronal IRE1α-deficient (IRE1α$^{SynCRe}$) mice at 6 weeks of age, with quantification shown in **(B)** (n = 6 mice per group). Data in (B) are presented as mean ± SEM and differences between IREα1$^{f/f}$ and IRE1α$^{SynCre}$ groups were analyzed using Student's *t* test. Significance levels are indicated as follows: *$p < 0.05$; **$p < 0.01$; ***$p < 0.001$: n.s., not significant.

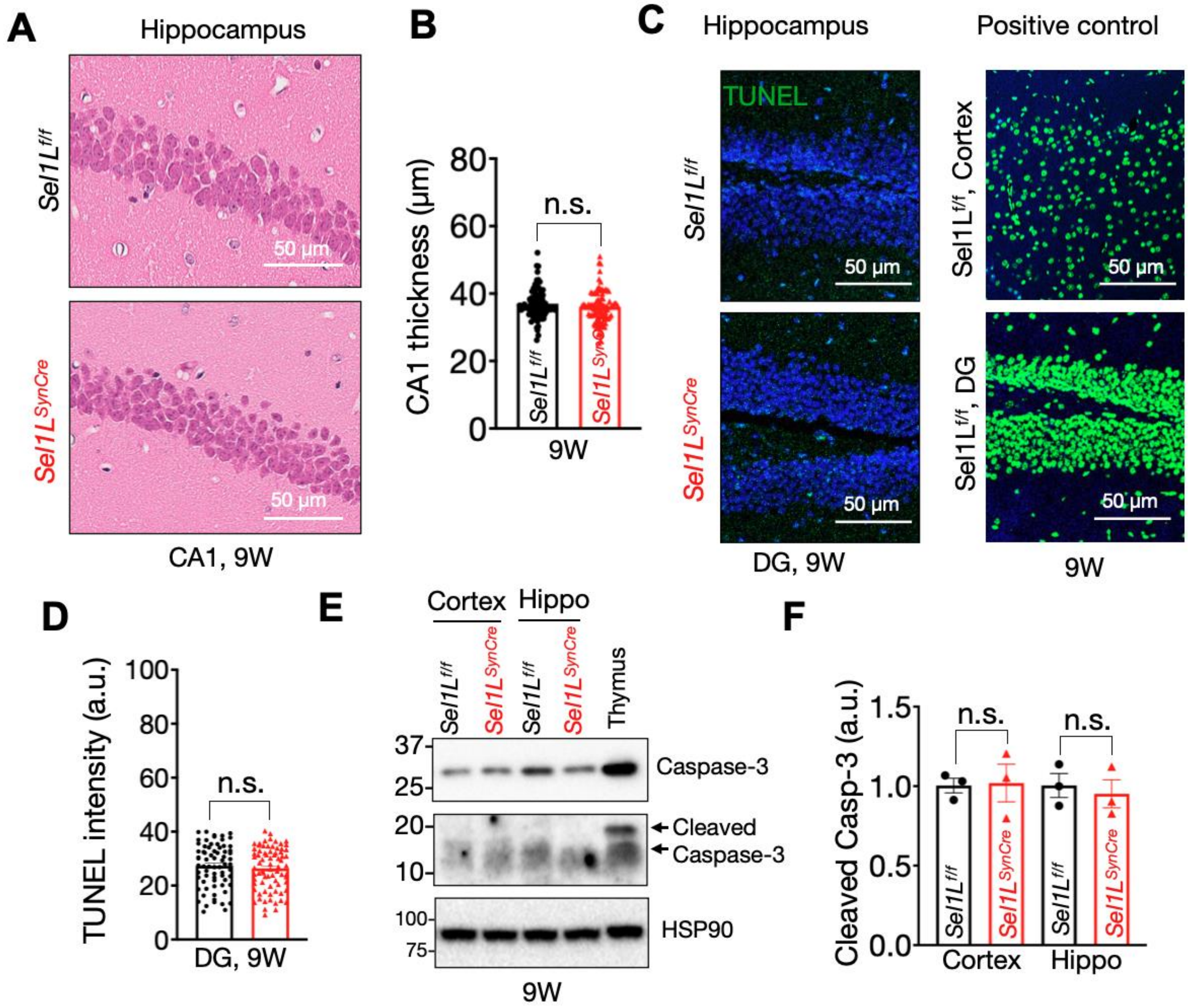


**Supplementary Figure 5. Neuronal SEL1L deficiency does not induce significant cell death.** (**A**) Hematoxylin & eosin staining of the hippocampus from *Sel1L*$^{f/f}$ and *Sel1L*$^{SynCre}$ mice at 9 weeks of age. Quantitation of the CA1 granular layer thickness is shown in (**B**) (50-60 measurements; n = 3 mice per group). (**C**) TUNEL assay for DNA fragmentation in hippocampal sections from Sel1L$^{f/f}$ and Sel1L$^{SynCre}$ mice at 9 weeks of age. A positive control with DNAse treatment showing cortex and dentate gyrus (DG) areas in a wild-type mice is shown on the right. Quantification of the average signal within a 50 x 50 µm$^2$ tissue area is shown in (**D**) (50-60 measurements; n = 3 mice per group). (**E**) Western blot analysis of cleaved caspase-3 in cortical and hippocampal tissue from Sel1L$^{f/f}$ and *Sel1L*$^{SynCre}$ mice at 9 weeks of age, with quantification shown in (**F**) (n = 3 mice per group). Data in B, D and F are presented as mean ± SEM and were analyzed using an unpaired two-tailed Student's *t* test (B and D) or two-way ANOVA followed by Tukey's multiple comparison test (F). Significance levels: *$p < 0.05$; **$p < 0.01$; ***$p < 0.001$: n.s., not significant.

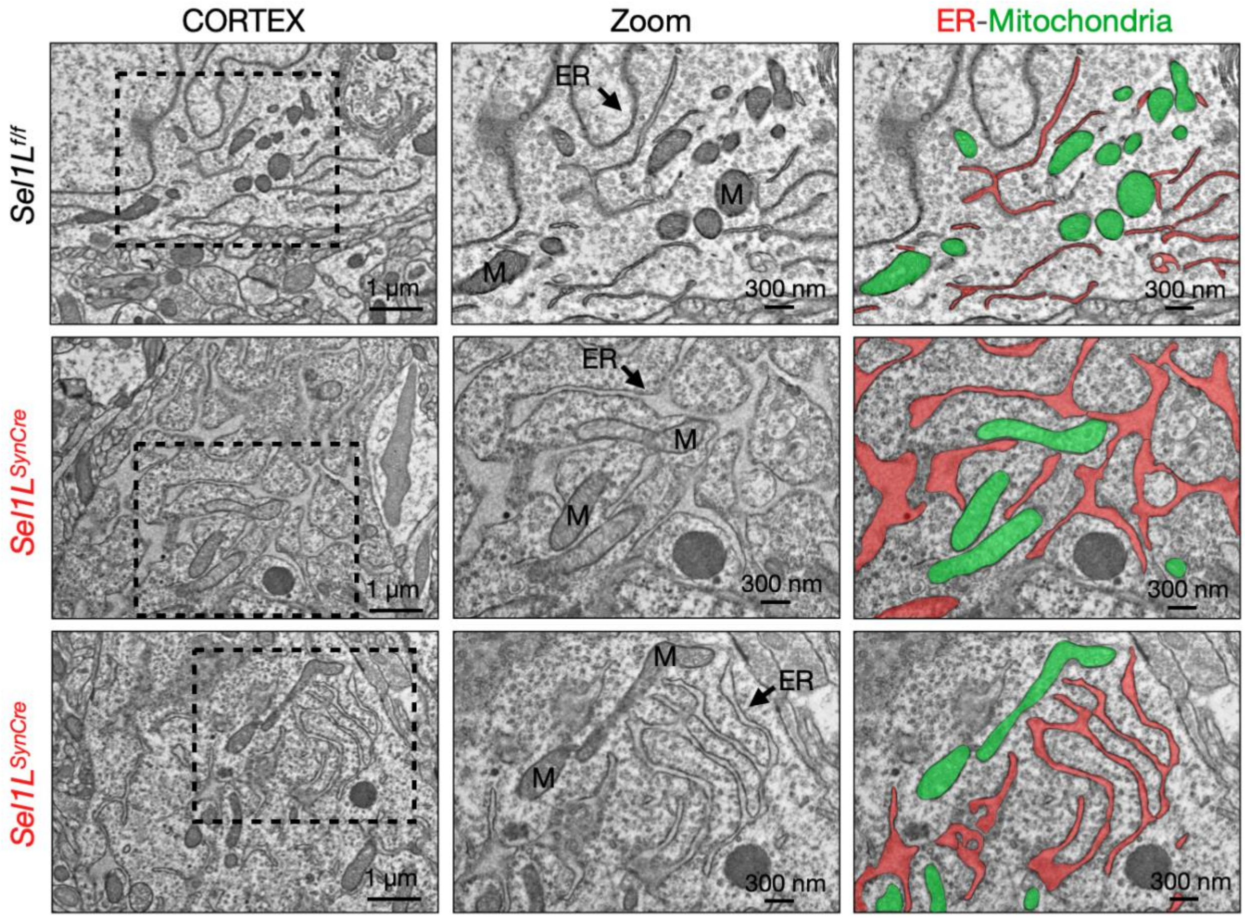


**Supplementary figure 6. ER expansion in cortical neurons deficient in Sel1L.**
Representative transmission electron microscopy (TEM) images from Sel1L$^{f/f}$ and Sel1L$^{SynCre}$ mice at 5 weeks of age. The endoplasmic reticulum (ER) and mitochondria (M) are highlighted in red and green, respectively. (n = 3 mice per group).

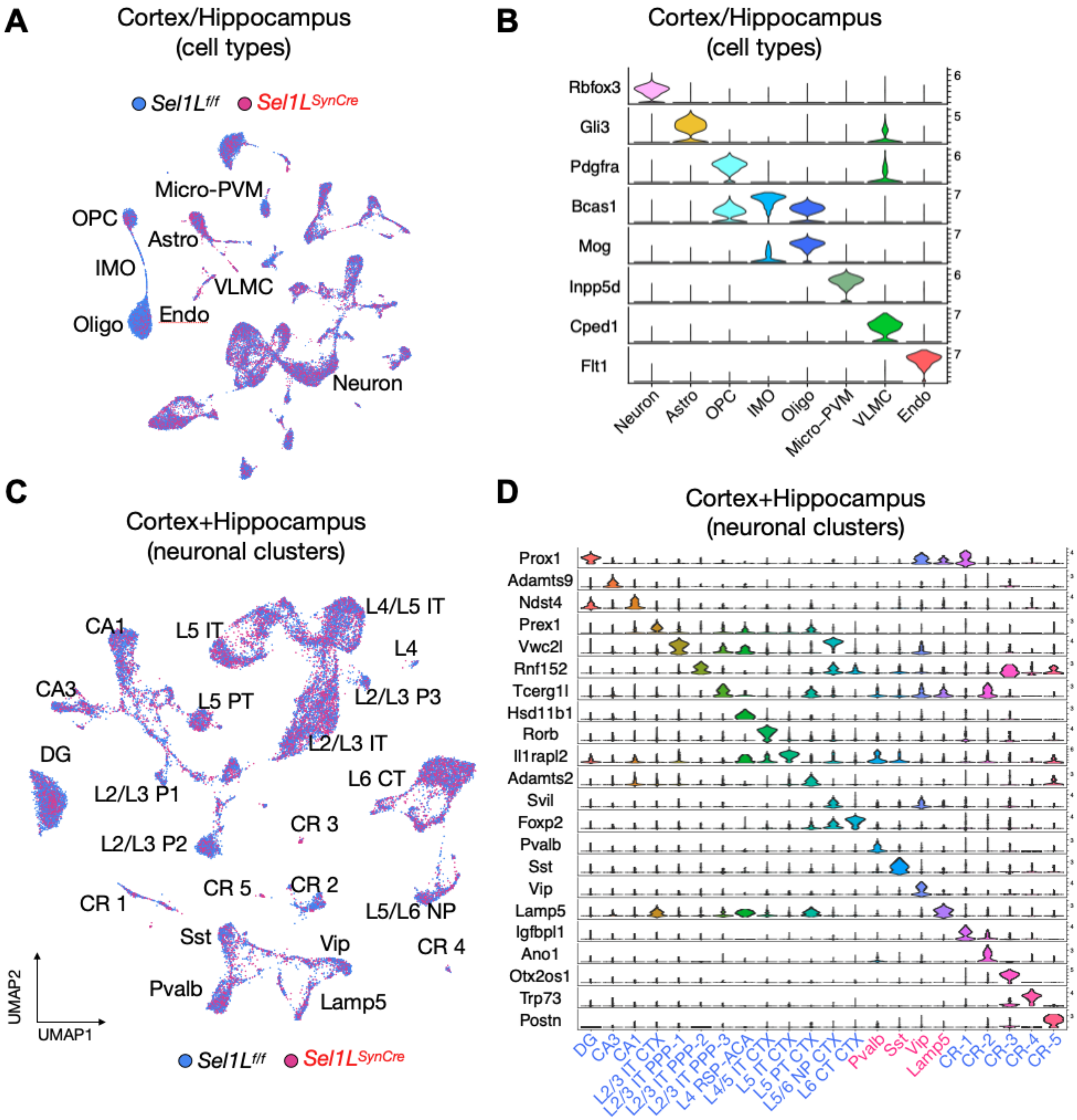


**Supplementary figure 7. Cell type separation and neuronal clustering in cortical and hippocampal tissue.** (**A**) UMAP plot showing the distribution of all cell types identified by single-nucleus RNA sequencing from cortical and hippocampal tissues of Sel1L$^{f/f}$ (blue) and Sel1L$^{SynCre}$ (red) mice at 5 weeks of age. (**B**) Violin plots showing the expression of marker genes used to identify different brain cell types. (**C**) Distribution of neuronal clusters identified in cortical and hippocampal tissues. (**D**) List of genes used to identify individual neuronal clusters present in cortical and hippocampal tissues. Clusters of excitatory and inhibitory neurons in panel D are shown in blue and red, respectively.

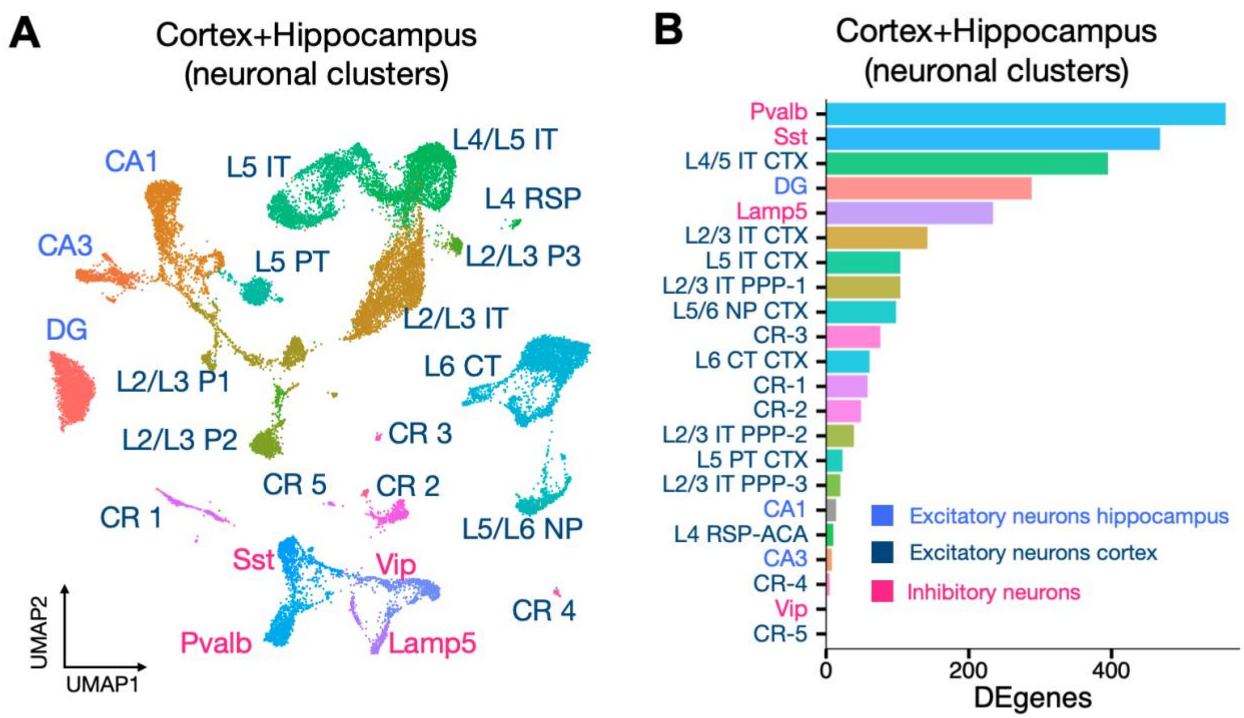


**Supplementary figure 8. Deferentially expressed genes in neuronal clusters from cortical and hippocampal tissues.** (**A**) Combined UMAP plot showing the distribution of neuronal clusters in cortical and hippocampal tissues of Sel1L$^{f/f}$ and Sel1L$^{SynCre}$ mice at 5 weeks of age. (**B**) Differentially expressed genes in neuronal clusters from Sel1L$^{SynCre}$ mice relative to *Sel1L*$^{SynCre}$ controls. cortical and hippocampal tissues of *Sel1L*$^{SynCre}$ mice relative to *Sel1L*$^{f/f}$ controls. Clusters exhibiting the most pronounced expression changes are listed at the top. Excitatory neuronal clusters in the cortex (dark blue) and hippocampus (light blue), as well as inhibitory neuronal clusters from both regions (red) are shown.

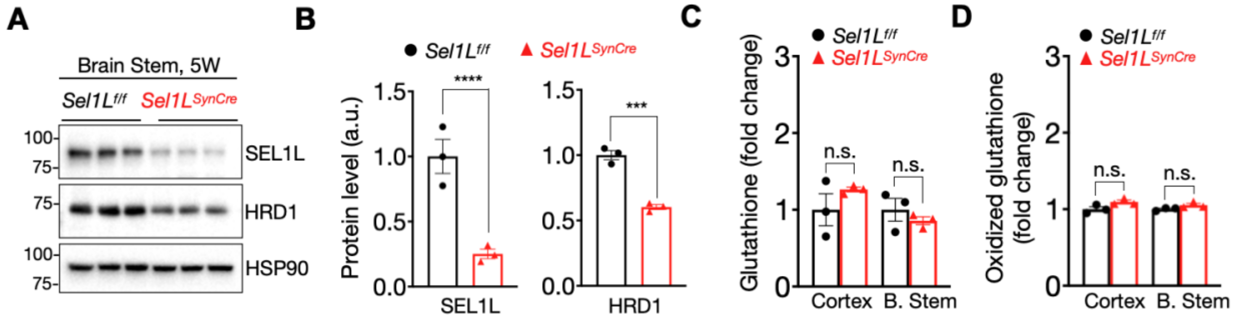


**Supplementary Figure 9. Activation of one-carbon metabolism pathways in the brain of Sel1L$^{SynCre}$ mice.** (**A**-**B**) Fold-change plots showing glutathione (**A**) and oxidized glutathione (**B**) metabolites from the cortex and brain stem of *Sel1L*$^{f/f}$ and *Sel1L*$^{SynCre}$ mice (n = 3 animals per brain region). (**C**) Western blot analysis of SEL1L and HRD1 proteins showing the efficient deletion of ERAD in brain stem at 5 weeks of age, with quantification shown in (**D**) (n = 3 mice per group). The data presented are shown as mean ± SEM and were analyzed using a Student's t test. Significance levels are indicated as follows: *p $<$ 0.05; **p $<$ 0.01; ***p $<$ 0.001.

### SUPPLEMENTAL VIDEOS

**Supplemental Video 1: Terminal-stage *Sel1L*$^{SynCre}$ mouse exhibiting pronounced tremor.** A 10-week-old *Sel1L*$^{SynCre}$ male mouse, alongside two *Sel1L*$^{f/f}$ control littermates, was recorded while consuming soft gel food in its home cage at the terminal stage. Note the persistent tremor observed during eating and grooming behavior.

**Supplemental video 2: Hindlimb clasping reflex in *Sel1L*$^{f/f}$ and *Sel1L*$^{SynCre}$ mice at 2 weeks of age.** The hindlimb clasping reflex was assessed via tail suspension in control (*Sel1L*$^{f/f}$) and neuron-specific *Sel1L* knockout (*Sel1L*$^{SynCre}$) mice at postnatal day 14. Healthy mice typically extend their hindlimbs outward when suspended by the tail. In contrast, *Sel1L*$^{SynCre}$ mice exhibit an abnormal clasping phenotype, retracting their hindlimbs toward the midline of the body.
